# A cytoplasmic chemoreceptor and Reactive Oxygen Species mediate bacterial chemotaxis to copper

**DOI:** 10.1101/2022.06.29.497744

**Authors:** Gwennaëlle Louis, Pauline Cherry, Catherine Michaux, Sophie Rahuel-Clermont, Marc Dieu, Françoise Tilquin, Laurens Maertens, Rob Van Houdt, Patricia Renard, Eric Perpete, Jean-Yves Matroule

**Affiliations:** Research Unit in Biology of Microorganisms (URBM), Department of Biology, Namur Research Institute for Life Sciences (NARILIS), University of Namur, Namur, Belgium; Laboratoire de Chimie Physique des Biomolécules, Namur Research Institute for Life Sciences (NARILIS) and Namur Institute of Structured Matter (NISM), University of Namur, Rue de Bruxelles, 61, 5000 Namur, Belgium; Université de Lorraine, CNRS, IMoPA, F-54000 Nancy, France; MaSUN, Mass Spectrometry Facility, University of Namur, Namur, Belgium; Microbiology Unit, Interdisciplinary Biosciences, Belgian Nuclear Research Centre (SCK CEN), 2400 Mol, Belgium

**Author notes:** Corresponding author. Mailing address: Research Unit in Biology of Microorganisms (URBM), 61 Rue de Bruxelles 5000 Namur, Belgium.

**Keywords:** Chemotaxis, bacteria, copper, reactive oxygen species (ROS), stress response

## Abstract

Chemotaxis is a widespread strategy used by unicellular and multicellular living organisms to maintain their fitness in stressful environments. We previously showed that bacteria can trigger a negative chemotactic response to a copper (Cu)-rich environment. Cu ions toxicity on bacterial cell physiology has been mainly linked to mismetallation events and ROS production, although the precise role of Cu-generated ROS remains largely debated.

Here, we found that the cytoplasmic Cu ions content mirrors variations of the extracellular Cu ions concentration and triggers a dose-dependent oxidative stress, which can be abrogated by superoxide dismutase and catalase overexpression. The inhibition of ROS production in the cytoplasm not only improves bacterial growth but also impedes Cu-chemotaxis, indicating that ROS derived from cytoplasmic Cu ions mediate the control of bacterial chemotaxis to Cu.

We also identified the Cu chemoreceptor McpR, which binds Cu ions with low affinity, suggesting a labile interaction. In addition, we demonstrate that the cysteine 75 and histidine 99 within the McpR sensor domain are key residues in Cu chemotaxis and Cu coordination. Finally, we discovered that *in vitro* both Cu(I) and Cu(II) ions modulate McpR conformation in a distinct manner. Overall, our study provides mechanistic insights on a redox-based control of Cu chemotaxis, indicating that the cellular redox status can play a key role in bacterial chemotaxis.

## Introduction

Cell motility is widely distributed across the three domains of life. For instance, in animals it is involved in fertilization, and in neutrophils recruitment to an inflammatory area. It also allows archaea and bacteria to seek nutrients in scarce environments, and to flee from toxic compounds while it supports to some extent host tropism in the frame of symbiotic relationships.

Most of the time, cell motility is driven by chemical gradients and is therefore named chemotaxis. In flagellated bacteria, positive (attraction) and negative (repulsion) chemotactic responses imply stochastic direction changes triggered by the inversion of flagellar rotation (1,2). This switch is caused by the binding of the phosphorylated response regulator, CheY∼P, to the flagellar motor. Spontaneous or phosphatase-mediated CheY∼P dephosphorylation by the CheZ phosphatase or by a member of the CheC/CheX/FliY phosphatase family, triggers CheY release from the flagellar motor (3–5). The release of CheY promotes a straight bacterial swimming pattern until CheY is phosphorylated again and triggers a new direction change. CheY phosphorylation relies on the histidine kinase CheA, which is recruited and autophosphorylated within a chemotaxis cluster composed of cytoplasmic or inner membrane-spanning methyl-accepting chemotaxis proteins (Mcp), also named chemoreceptors (6). Chemoreceptors are classically composed of a N-terminal sensing domain, trans-membrane (TM) domains (which are absent in cytoplasmic chemoreceptors), a HAMP domain and the C-terminal (Mcp) signaling domain. Depending on their cellular topology, chemoreceptors monitor chemical variations in the cytosol or in the periplasm *via* their N-terminal sensing domain. Upon substrate binding to their sensing domain, chemoreceptors undergo a conformational change leading to the increase (repellent response) or the decrease (attractant response) of CheA histidine kinase activity (7,8). In *E. coli* Tar and Tsr chemoreceptors, this conformational change consists in a 2 Å shift of one of the TM helices (TM2), causing a “piston-like” movement in the direction of the membrane that impacts the HAMP domain (9). The methylation of the Mcps by the CheR methyltransferase results in an increase of CheA activity and their demethylation by the CheA-activated CheB methylesterase leads to the decrease of CheA activity (9). This adaptation response sets back the Mcps in a pre-stimulus state.

Copper (Cu) has been used for centuries as an antimicrobial and antifouling strategy in medicine and in various anthropogenic activities including agriculture and industry. Cu toxicity in solution mainly results from its high redox potential causing the displacement of native metallic ions from metalloproteins, and the disruption of Fe-S clusters (10). Cu ability to generate reactive oxygen species (ROS) has been shown *in vitro* (11) but the role of these ROS *in vivo* yet remains unclear. In a previous study, we identified an original bimodal strategy to cope with Cu stress where the flagellated morphotype (swarmer cell) of *Caulobacter crescentus* triggers a prompt negative chemotaxis from the Cu source (12). In addition to the negative Cu-chemotaxis, *C. crescentus* exhibits a positive chemotactic behavior toward xylose (13) and O_2_ (14) but no dedicated Mcp could be identified so far.

*C. crescentus* genome harbors two chemotaxis operons. The major chemotaxis operon encodes McpA and McpB, CheYI and CheYII, CheAI, one CheW, CheRI, CheBI and the CheY-like CleA. This operon also contains the *cheD*, *cheU* and *cheE* genes, which have been hardly described. CheYII and CheBI are likely the major CheY and CheB in *C. crescentus* chemotaxis (15,16). The alternate chemotaxis operon encodes McpG and McpK, two CheY, CheAII, one CheW, CheBII, and CheRII. The other *cheY* genes and the third *cheW* are dispersed within the genome. Only the first chemotaxis operon seems to be essential for chemotaxis while both operons are involved in biofilm formation and holdfast production, by regulating the expression of the *hfiA* holdfast inhibitor (16,17).

No CheZ homolog has been found in *C. crescentus* genome but the multiple CheY are proposed to act as phosphate sinks to allow the termination process, such as in *Rhizobium meliloti* (*18*). Five of the CheY have been redefined as Cle proteins (CheY-like c-di-GMP effectors) and renamed CleA to CleE. CleA and CleD could compete with CheYII to prevent the rotational switch of the flagellum and to promote straight run (15). The *C. crescentus* MCPs array exhibits a higher-ordered hexagonal structure constituted of approximatively one to two thousand MCPs per array (19). Among the *C. crescentus* MCPs, only the inner membrane McpA and the cytoplasmic McpB encoded by the major chemotaxis operon were described as polar chemoreceptors, although their exact function remains unknown (20).

In the present study, we aimed to decipher the sensing mechanism underlying *C. crescentus* chemotaxis to a toxic Cu-rich environment. We provide evidence that an increased intracellular Cu concentration triggers a dose-dependent production of reactive oxygen species (ROS), which is a prerequisite to Cu chemotaxis. We also demonstrate that the redox status of the Cu ions bound to the sensor domain of the newly identified McpR modulates McpR conformation *in vitro*. This original redox sensing mechanism may ensure a continuous monitoring of the intracellular Cu status, providing a real-time control of the chemotactic response.

## Results

To further dissect the molecular mechanisms underlying the negative Cu-chemotaxis of *Caulobacter crescentus* swarmer (SW) cells (12), we first sought to identify the chemoreceptor involved in Cu chemotaxis. The genome of *C. crescentus* is predicted to encode 19 chemoreceptors (named McpA to McpS) based on the presence of an Mcp domain binding CheA/CheW (21). No canonical Cu ions-binding site, including the CXXC motif or the βαββαβ fold (22), could be found by running manual and JPred4-based analyses of the 19 chemoreceptor protein sequences (23). Therefore, we proceeded to single in-frame deletions of the 19 *mcp* genes, and their respective chemotactic behavior to Cu ions was assessed by using the previously designed Live Chemotaxis Imaging (LCI) (12). Most of the Δ*mcp* mutants displayed a WT-like behavior with a flight percentage after 25 min varying between 15% and 51%. Only Δ*mcpA*, Δ*mcpB* and Δ*mcpR* mutants showed a significant decrease of Cu-chemotaxis relative to the WT strain (**Fig. 1A**), suggesting that these chemoreceptors might be involved in Cu-chemotaxis. The Δ*mcpA*, Δ*mcpB* and Δ*mcpR* mutants could be complemented by the expression of an ectopic version of *mcpA*, *mcpB* and *mcpR* on a low copy plasmid, respectively, ruling out any polar effect of the deletions (**Fig. S1**). Owing to the position of the *mcpA* gene within the major chemotaxis cluster (16), McpA is likely involved in chemotaxis to a less specific extent. Consistent with this hypothesis, the Δ*mcpA* mutant displays a reduced motility halo in a Cu ions-free swarming assay, while the Δ*mcpB* and Δ*mcpR* mutants exhibit a WT-like chemotaxis profile in the same condition (**Fig. S2A**). The presence of the terminal sequence of the CheB/CheR docking site in McpB (absent from McpR) led us to the idea that McpB might be necessary for the adaptive response as in *E. coli* (24), which will not be considered in this study. Therefore, we decided to determine how McpR modulates Cu-chemotaxis.

**Figure 1:**
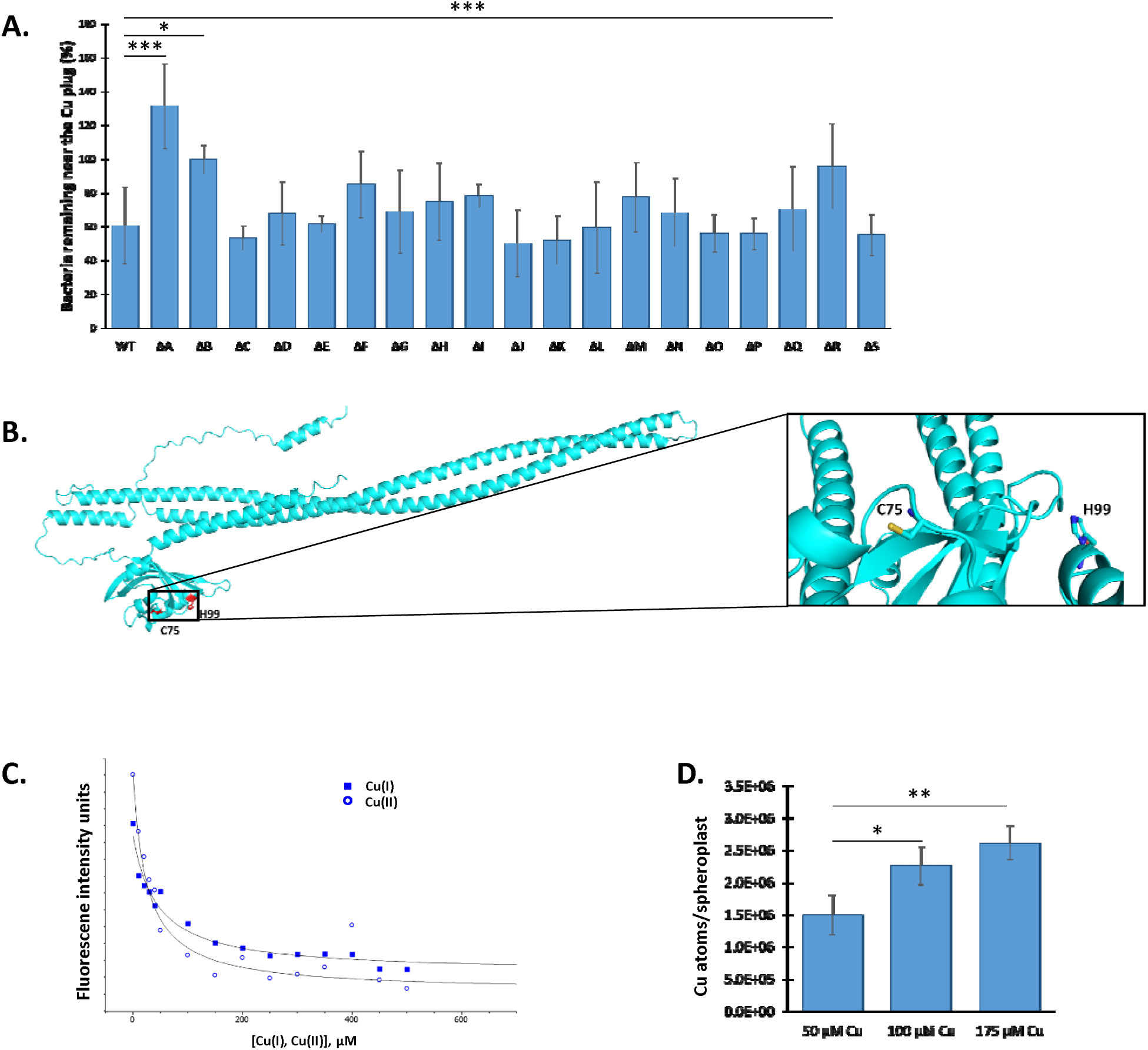
McpR is involved in Cu chemotaxis and Cu binding. **A.** Percentage of WT and Δ*mcp (mcpA* to *mcpS)* SW cells remaining in the vicinity of the Cu plug after 25 min. **B.** 3D structure prediction of McpR of McpR by AlphaFold. H99 and C75 residues are highlighted. **C.** Representative McpR titration by Cu(I) (squares) or Cu(II) (circles) monitored by protein intrinsic fluorescence intensity quenching, fitted to a single-site hyperbolic model by non-linear regression analysis (solid line). **D.** Number of Cu atoms per spheroplast from SW cells exposed to various Cu concentrations for 5 min. Mean +/- s.d., at least 3 biological replicates. *p-values* were calculated using an ANOVA (* *p* < 0.05, ** *p* < 0.01, *** *p* < 0.001) (**Table S1**).

The ability of the Δ*mcpR* mutant to form a motility halo in a Cu ions-free swarming assay (**Fig. S2A**) and to sustain negative chemotaxis to other chemicals, such as zinc (Zn) ions (**Fig. S2B**) indicate that the loss of Cu-chemotaxis in the Δ*mcpR* mutant does not result from a loss of motility and/or a disruption of the general chemotactic machinery.

McpR harbors the canonical domain organization found in chemoreceptors including a N-terminal sensing domain followed a HAMP domain and a C-terminal Mcp domain. McpR sequence analysis using the TMHMM 2.0 (25) and SignalIP 6.0 (26) tools did not reveal any McpR transmembrane domain and signal peptide respectively, arguing for a cytoplasmic localization for McpR, and therefore a sensing of the cytoplasmic Cu content.

Chemoreceptors can sense attractants or repellents in a direct or indirect manner *via* their sensing domain (27). The 3D structure prediction of McpR by Alphafold2 (28,29) highlights a cysteine (C75) in the vicinity of a histidine (H99) in the McpR sensing domain (**Fig. 1B)**. Cysteine and histidine residues have often been described as key residues in Cu coordination (30), suggesting that McpR may bind Cu ions directly.

Therefore, we used intrinsic fluorescence spectroscopy to assess the direct binding of Cu to the purified McpR. Upon the addition of either Cu(I) or Cu(II) ions, the intrinsic McpR fluorescence at 346 nm is quenched in a dose-dependent manner, reflecting a modification of the McpR aromatic residues environment (**Fig. 1C**). The dissociation constants (*K*d) (see Materials and Methods for calculation method) of 48 µM +/- 10 µM for Cu(I) ions and 37 µM +/- 9 µM for Cu(II) ions attest to a similar affinity of both ionic forms for McpR. However, the micromolar range of Cu(I) ions and Cu(II) ions *K*ds exceeds from several logs the *K*d of well-known cuproproteins (31,32), arguing for a rather labile interaction between McpR and Cu. Nevertheless, McpR seems to preferentially bind Cu ions since Zn, cadmium (Cd), manganese (Mn) and nickel (Ni) ions had no effect on purified McpR intrinsic fluorescence (**Fig. S3**).

To determine whether the cytoplasmic Cu ions content could reflect variations of the extracellular Cu ions concentration when the SW cell swims away from a Cu ions source, we measured by ICP-OES the Cu ions content in cytoplasmic fractions from SW cells subjected to increasing Cu ions concentrations. Although the measurement cannot discriminate between free and complexed Cu ions, Cu(I) and Cu(II), we measured a dose-dependent accumulation of Cu ions in the cytoplasm (**Fig. 1D**). This observation is in accordance with previous data from our lab, showing that an artificial decrease of Cu ions content *via* the overexpression of the PcoB efflux pump impeded Cu-chemotaxis (12). Together, these data suggest that the sensing of cytoplasmic Cu ions by McpR could be used as a proxy by the SW cells to mirror variations of the concentration of Cu ions in *C. crescentus* environment to control Cu-chemotaxis.

The transient maintenance of a high intracellular Cu ions content in the SW cells may potentially lead to an oxidative stress *via* the production of superoxide anion (O_2_ ^•**-**^), hydrogen peroxide (H O) and hydroxyl radical (OH^•^) (**Fig. S4A**). In line with this idea, the relative fluorescence intensity of the ROS sensitive turn-on CellROX Deep Red probe is significantly increased in SW cells exposed to Cu ions, and H_2_O_2_, the latter being used as a positive control (**Fig. 2A**). Accordingly, the fluorescence of the cytoplasmic turn-off rxYFP redox biosensor is partially quenched under the same conditions (**Fig. 2B**), mirroring an imbalance of the cytoplasmic GSH-GSSG pool upon oxidative stress (**Fig. S4B**). These findings are reinforced by the differential expression of genes involved in oxidative stress response upon Cu stress, including those encoding the catalase KatG and the superoxide dismutases SodA and SodB (33). As observed for the cytoplasmic Cu ions content (**Fig. 1D**), the amount of ROS is increased when the SW cells are exposed to increasing extracellular Cu ions concentrations, indicating that the cytoplasmic ROS level mirrors the cytoplasmic Cu ions content (**Fig. 2C**).

**Figure 2:**
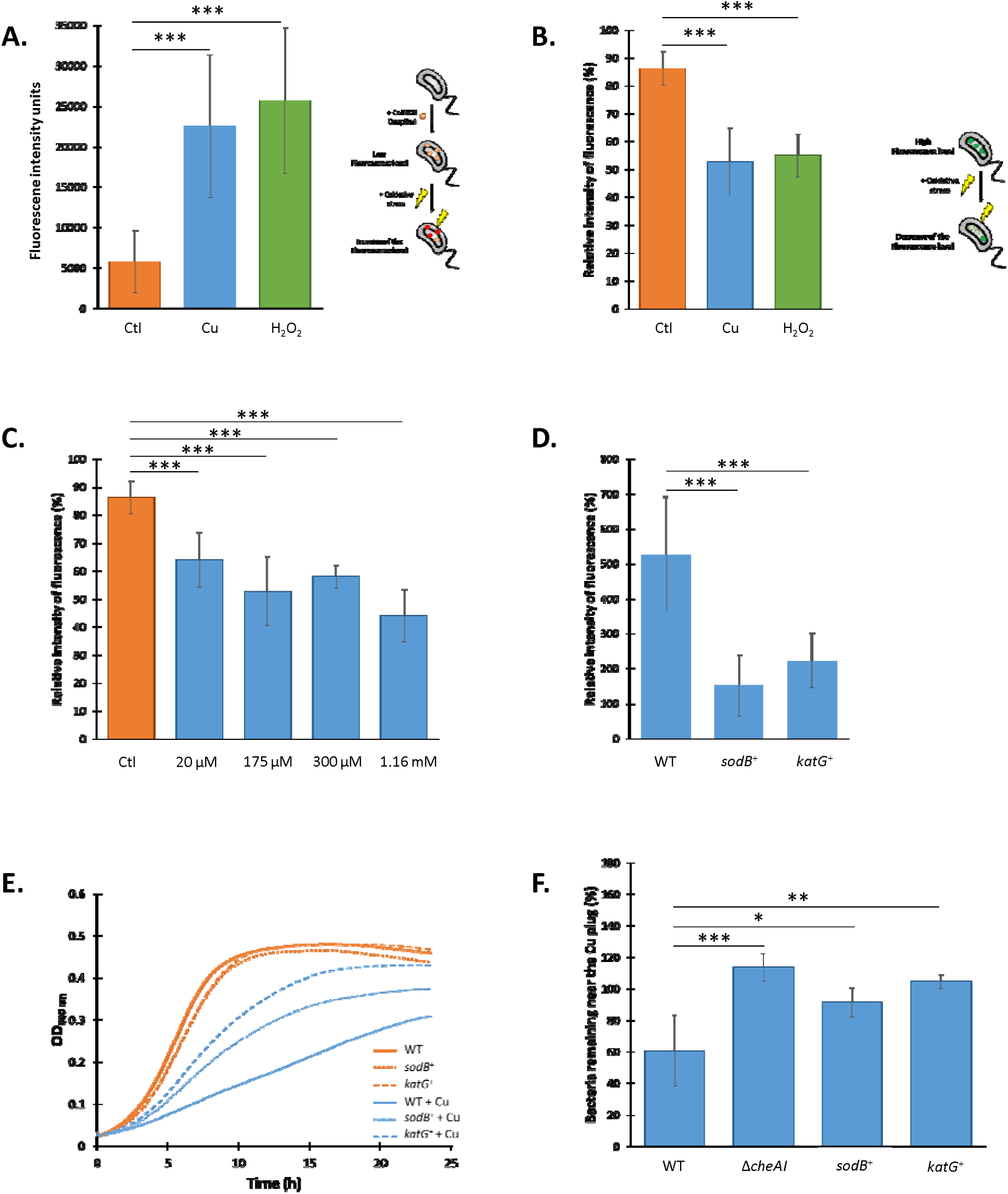
Cu-induced ROS sustain Cu chemotaxis. **A.** Raw fluorescence of SW cells incubated with 5 µM CellROX Deep Red probe and exposed to 175 µM Cu or 400 µM H_2_O_2_. **B.** Relative fluorescence intensity of SW cells expressing the rxYFP biosensor and exposed to 175 µM Cu or 400 µM H_2_O_2_ for 20 min. **C.** Relative fluorescence intensity of SW cells expressing the rxYFP biosensor and exposed to various Cu concentrations for 20 min. **D.** Relative fluorescence intensity of WT SW cells and SW cells overexpressing SodB (*sodB+*) and KatG (*katG*+) incubated with 5 µM CellROX Deep Red probe and exposed to 175 µM Cu. **E.** Growth profiles at OD_660 nm_ of WT, *sodB+* and *katG*+ strains in PYE (orange) and exposed to 175 µM Cu (blue). **F.** Percentage of WT, Δ*cheAI*, *sodB+* and *katG+* SW cells in the vicinity of the Cu plug after 25 min. Mean +/- s.d., at least 3 biological replicates, except for 300 µM Cu (2 biological replicates). *p-values* were calculated using an ANOVA when appropriate (* *p* < 0.05, ** *p* < 0.01, *** *p* < 0.001) (**Table S1**).

We envisioned that boosting the antioxidant defense system would limit ROS production by Cu ions. Therefore, we ectopically expressed the *sodB* or the *katG* genes in the WT background from a low copy plasmid under the control of the strong and constitutive *lac* promoter. The resulting *sodB*^+^ and *katG^+^*strains prevent the increase of the CellROX Deep Red fluorescence upon SW cells exposure to Cu ions, reflecting a potentiation of antioxidant defenses (**Fig. 2D**). In support of this data, the growth of these two optimized *sodB*^+^ and *katG^+^* strains is improved under moderate Cu stress (**Fig. 2E**), arguing for a role of O_2_ ^•**-**^ and H O in Cu toxicity.

Interestingly, the *sodB*^+^ and *katG^+^*strains are impeded in Cu-chemotaxis similarly to the Δ*cheAI* mutant (**Fig. 2F**, **SI Appendix and Fig. S5**), indicating that the ROS resulting from the accumulation of cellular Cu ions play a role in Cu-chemotaxis. It is worth mentioning that the loss of Cu-chemotaxis in the Δ*mcpR* mutant is not due to a loss of ROS production under Cu stress since the Δ*mcpR* mutant exhibits a similar decrease of the rxYFP fluorescence to the WT strain upon Cu ions exposure (**Fig. S6**). These data support a major role of the ROS resulting from Cu ions accumulation in Cu-chemotaxis and indicate that Cu ions binding to McpR is solely not sufficient to sustain Cu-chemotaxis.

Owing to the oxidative conditions resulting from Cu ions accumulation in the cytosol (**Fig. 2A-C)**, we sought to assess by LC/MS the oxidation status of McpR *in vitro* after incubating the purified McpR with Cu(II) ions or Cu(I) ions +/- H_2_O_2_. Only the Cu(I) ions + H_2_O_2_ condition, and to a lesser extent Cu(I) ions, led to a significant oxidation of the McpR sensor domain (residues 1-161), and more specifically of the H99 residue (**Fig. 3A, left panel**). In this condition, about 30% of the H99 residues acquired an additional oxygen atom, most likely yielding 2-oxoHis. Metal catalyzed oxidation (MCO) of histidines via the generation of OH^•^ has been previously described in the *Bacillus subtilis* PerR repressor (34), in yeast SOD1 (35), and in amyloid beta (Aβ) peptide (36). The exact role of this modification remains unknown and no redox cycling involving 2-oxoHis has been described so far, questioning the relevance of H99 oxidation in Cu/ROS sensing. MCO mostly targets the metal-binding residues in a caged process (37), suggesting that H99 may coordinate Cu. To test this hypothesis, we purified a McpR_H99A_ mutant protein and calculated the *K*d values of the McpR_H99A_/Cu(I) ions and McpR_H99A_/Cu(II) ions complexes (**Fig. 3B, Fig. S7A for the fitting curves**). H99A mutation triggers a 9.4- and 5.8-fold increase of Cu(I) ions and Cu(II) ions *K*d, respectively, relative to McpR, suggesting a key role of H99 in Cu(I) ions and Cu(II) ions coordination. The important variability associated with the McpR_H99A_ *K*d values likely reflects a potential instability of this mutant in solution upon Cu(I) ions and Cu(II) ions exposure, resulting in the presence of several subpopulations of McpR_H99A._

**Figure 3:**
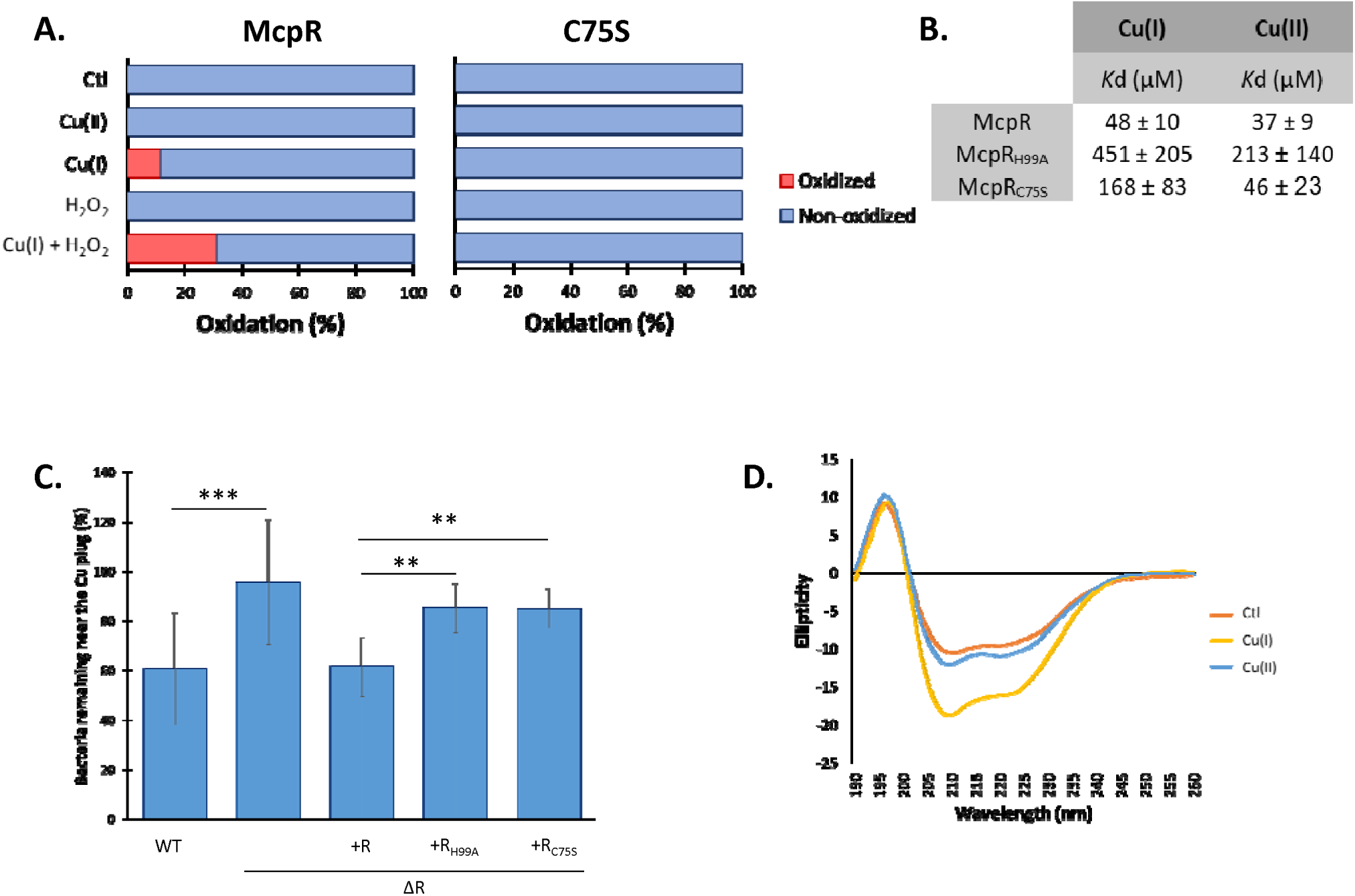
Cu cycling modulates its coordination by McpR. **A.** Percentage of oxidation of the H99 residue from purified McpR (left panel) and McpR_C75S_ (right panel) incubated with Cu(II), Cu(I), H_2_O_2_ and Cu(I)+H_2_O_2_ at a 1:100 ratio. **B.** Measure, by intrinsic fluorescence, of the *K*d values of purified McpR, McpR_H99A_ and McpR_C75S_ (1.79 µM) incubated with 0 to 500 µM Cu(I) and Cu(II). Mean +/- s.d. **C.** Percentage of WT, Δ*mcpR* and Δ*mcpR* overexpressing McpR, McpR_H99A_ or McpR_C75S_ SW cells remaining in the vicinity of the Cu plug after 25 min. Mean +/- s.d., at least 3 biological replicates. *p-values* were calculated using an ANOVA (* *p* < 0.05, ** *p* < 0.01,*** *p* < 0.001) (**Table S1**). **D.** Circular Dichroism spectra of purified McpR (8.95 µM) incubated with Cu(I) and Cu(II) at a 1:100 ratio.

To assess the potential implication of the neighboring C75 residue in Cu ions coordination, we purified the McpR_C75S_ mutant protein and calculated the *K*d values of the McpR_C75S_/Cu(I) ions and McpR_C75S_/Cu(II) ions complexes. As for the H99A mutation, the C75S mutation also impeded Cu(I) ions binding as attested by a 3.5-fold *K*d increase. Cu(II) ions binding to the McpR_C75S_ mutant remained unchanged (**Fig. 3B, Fig. S7B for the fitting curves**), suggesting that McpR C75 is required to coordinate Cu(I) ions but not Cu(II) ions. Consistent with this observation, *in vitro* H99 oxidation by Cu(I) ions +/- H_2_O_2_ is fully abolished in the McpR_C75S_ variant (**Fig. 3A, right panel)**. The *K*d values calculated for the McpR_C75S_ and McpR_H99A_ variants are in agreement with the known selective binding of Cu(I) and Cu(II) ions to cysteine and histidine residues (30,38).

To determine whether Cu coordination by McpR H99 and C75 residues is essential to Cu-chemotaxis, we decided to express an ectopic *mcpR_H99A_*or *mcpR_C75S_* allele in the Δ*mcpR* mutant and to monitor the chemotactic behaviour of both strains to Cu ions. None of the mutant allele was able to complement the Cu-chemotaxis defect of the Δ*mcpR* mutant (**Fig. 3C)**. One could argue that the McpR_H99A_ and McpR_C75S_ variants are unstable *in vivo*, explaining the Δ*mcpR* mutant phenotype. In accordance with the pleiotropic sensing ability of some Mcps, we discovered that McpR is also playing a key role in cadmium (Cd)-chemotaxis (**Fig. S8**). Interestingly, the *mcpR_H99A_*and *mcpR_C75S_* mutant strains were not impeded in Cd-chemotaxis (**Fig. S8**). This latter observation attests the integrity of the McpR_H99A_ and McpR_C75S_ variants *in vivo* and suggests that Cd sensing by McpR does not rely on the H99 and C75 residues.

These data led us to the idea that Cu(I) ions coordination by McpR H99 and C75 residues is decisive for McpR-mediated Cu chemotaxis. Given the structural changes elicited by the binding of a ligand to a cognate chemoreceptor (39), we envisioned that Cu(I) ions binding to McpR could modulate its conformation. To test this idea, Far-UV circular dichroism (CD) spectroscopy was used to monitor the impact of Cu(I) ions and Cu(II) ions on McpR secondary structure *in vitro*. Accordingly, Cu(I) ions has a more significant impact than Cu(II) ions on McpR secondary structure (**Fig. 3D)**, increasing the α-helix percentage from 23% to 44%.

Based on our findings, we propose that Cu exposure triggers a dose-dependent Cu accumulation within the cytoplasm triggering a dose-dependent production of ROS that will sustain Cu chemotaxis. The increase in cytoplasmic Cu ions concentration will likely favor a rather labile Cu ions binding to the McpR sensor domain. Cu(I) ions coordination by McpR C75 and H99 residues triggers McpR conformational change. Interestingly, Cu(II) ions seem to preferentially bind McpR H99 residue and does not modify the apo form-like conformation. These observations suggest that a redox cycling of the McpR-bound Cu atom could modulate McpR conformation (**Fig. 4**).

**Figure 4:**
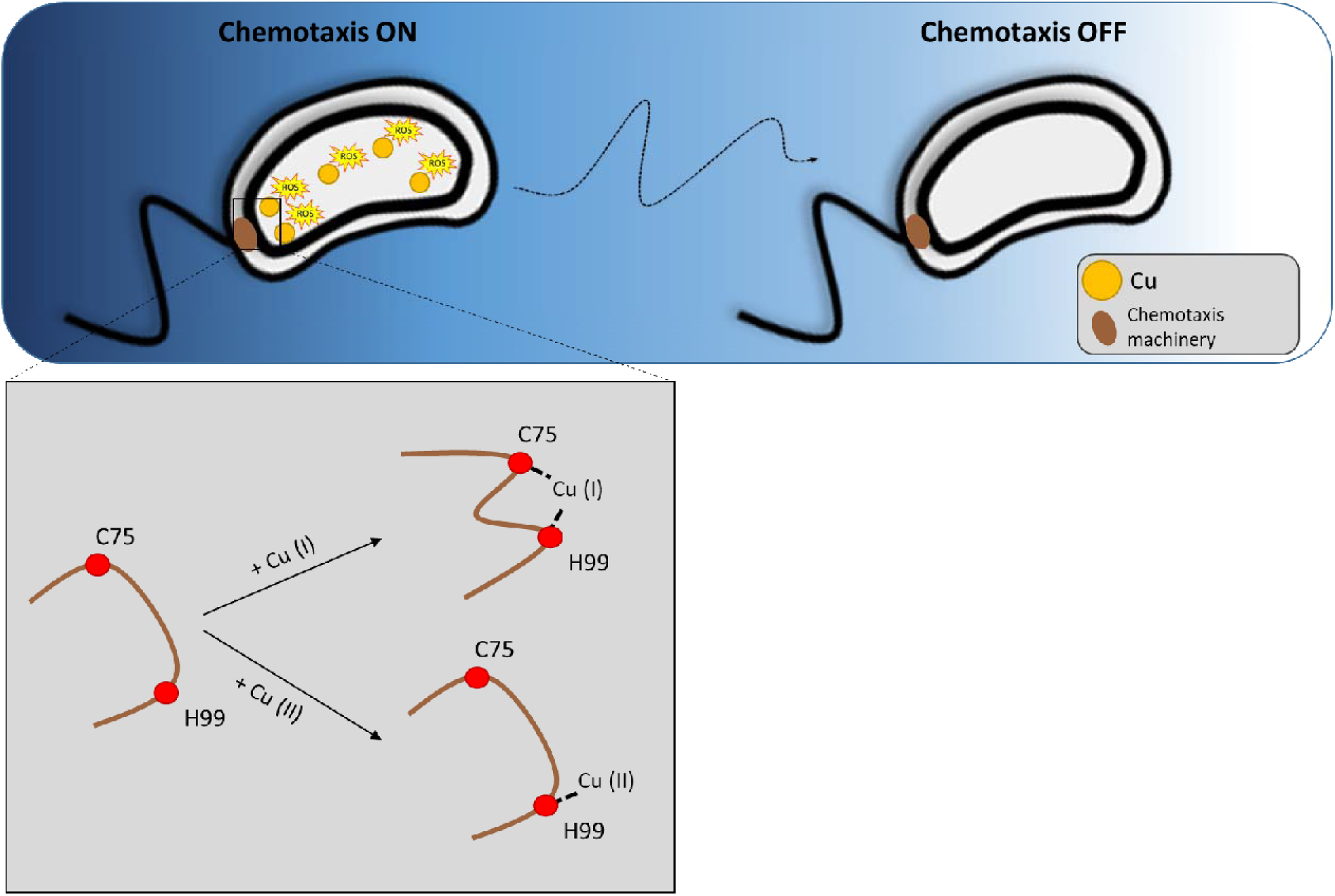
Hypothetical model of Cu sensing by McpR. SW cell exposure to Cu increases the cytoplasmic Cu concentration, leading to the generation of ROS (left). In addition, high cytoplasmic Cu content favors Cu(I) coordination by the C75 and H99 residues within the cytoplasmic McpR, triggering a conformational change of the chemoreceptor. Cu(II) binding to H99 residue has no impact on McpR apo-like conformation, suggesting a redox-dependent modulation of McpR conformation. This process would be maintained along the extracellular Cu gradient until the SW cell reaches a Cu-poor environment (right). Under these conditions, the cytoplasmic Cu concentration together with the ROS level would return to their basal level and the labile Cu binding to McpR would be hindered, bringing McpR to its apo form conformation.

## Discussion

The role of ROS in Cu toxicity and Cu stress response remains largely unknown in bacteria. We sought to address this gap in the frame of negative chemotaxis to toxic Cu in the dimorphic bacterial model *C. crescentus*.

Our study identified the first Cu sensing chemoreceptor hereafter named McpR. McpR is devoid of any transmembrane domains or signal peptide and is therefore predicted to localize in the cytoplasm. Cytoplasmic chemoreceptors have been shown to monitor the cellular metabolic state (40), whereas inner membrane chemoreceptors usually sense the periplasmic environment, as a proxy for extracellular changes. The cytoplasmic Cu content varies in a dose-dependent manner when the SW cells are exposed to Cu, suggesting that the sensing of cytoplasmic Cu could provide a sensitive monitoring of subtle variations of Cu concentrations when the SW cell swims down a Cu gradient.

In a previous study, we showed that the maintenance of a high cellular Cu concentration is critical for Cu chemotaxis (12). Consistent with this observation, we provide evidence that McpR binds Cu with low affinity. Although aspecific Cu binding to non-cognate residues cannot be ruled out, the significant difference in *Kd* values measured between the WT McpR and the McpR_H99A_ and McpR_C75S_ variants argue for a Cu binding to these residues. Considering that Cu binding to McpR is a prequisite for Cu chemotaxis, one could propose that when the SW cell is sufficiently far from the Cu source, the resulting low cytoplasmic Cu content impedes Cu binding to McpR and in turn the chemotactic response.

We also demonstrate that the cellular Cu accumulation triggers a dose-dependent production of ROS. This dose dependency in ROS production argues that the increase of extracellular Cu ions content triggers an increase of the cytoplasmic Cu ions labile pool, which is likely involved in the labile Cu binding to McpR. ROS production can be prevented by the overexpression of SodB superoxide dismutase and KatG catalase, suggesting that at least O **^.-^** and H O are generated under Cu stress. The contribution of the ROS derived from cellular Cu ions in Cu toxicity is still largely debated (41,42). Here, we demonstrate that ROS derived from cellular Cu ions not only impede cell growth to some extent, but also play a key role in Cu-chemotaxis. The exact target of ROS in Cu-chemotaxis was not identified in this study, but one could hypothesize that O **^.-^** is rapidly converted into H_2_O_2_. The latter would feed a local Fenton-like reaction catalyzed by the Cu(I) atom coordinated by McpR C75 and McpR H99 residues. This would lead to a local OH**^.^** production and to the oxidation of McpR-bound Cu(I) into McpR-bound Cu(II). Owing to the reducing cytoplasmic environment, McpR-bound Cu(II) could in turn be reduced by cytoplasmic electron donors such as GSH, NADH and thioredoxin to restore the Cu(I) ionic form. Thereby, the McpR-bound Cu ion would undergo a redox cycling involving the Cu(I) and Cu(II) ionic forms, which would be maintained as long as H_2_O_2_ is produced by cytoplasmic Cu ions. Cu-generated ROS would therefore act as second messengers in a bacterial stress response.

The modulation of the CheA-CheY two-component system results from a conformational change of the chemoreceptor occurring upon ligand binding (9). We demonstrated here that McpR conformation is modulated by the ionic status of the Cu atom bound to McpR. It should be emphasized that chemoreceptors are usually organized in trimers of dimers and assembled in large arrays in cells (43). Therefore, it is possible that Cu coordination by the McpR C75 and H99 residues within the McpR array implies two McpR monomers from one or two distinct dimers. Nevertheless, McpR conformational change likely regulates the downstream effectors such as CheAI and the resulting chemotactic response.

The diversity of chemoreceptors produced by a single bacterial species varies from 5 to 60 chemoreceptors and is often correlated to its lifestyle (40). Nevertheless, chemoreceptors are not always specific to one single substrate and can also perform indirect sensing *via* a substrate transporter (27). For instance, the *E. coli* Tar chemoreceptor directly senses aspartate and performs an indirect sensing of maltose through a periplasmic maltose binding protein (44).

The extrapolation of our findings to other motile bacterial species based on sequences analyses is complicated by (i) the large diversity of Cu coordination mechanisms and thereby the low predictability of Cu-binding sites by *in silico* analyses and (ii) the low conservation of the Mcp sensor domains. Nevertheless, it is very likely that the use of ROS as second messengers is widely conserved in bacterial response to Cu.

## Experimental procedures

### Bacterial strains, plasmids and growth conditions

*Caulobacter crescentus* (NA1000 strain) was grown at 30°C in Peptone Yeast Extract (PYE) medium (45) with 5 μg/ml kanamycin, 1 µg/ml chloramphenicol and/or an appropriate concentration of CuSO_4_.5H_2_O when required. Cultures in exponential phase were used for all experiments. Strains, plasmids and primers used in this study are listed in Table S2, and the strategies for their construction are available upon request. SW cells isolation was achieved according to (46). Briefly, the bacteria were collected at an O.D.^660 nm^ of 0.4 and centrifuged in Ludox LS colloidal silica gradient. After the centrifugation, SW are separated from the other cellular forms (ST and PD) based on a density difference. The SW cells were collected and washed several times with PBS prior the experiments.

### Growth curve measurements

*C. crescentus* exponential phase cultures were diluted in PYE medium to a final O.D._660 nm_ of 0.05 and inoculated in 96-well plates with appropriate CuSO_4_ concentration when required. Bacteria were then grown for 24 h at 30°C under continuous shaking in an Epoch 2 Microplate Spectrophotometer from BioTek (Santa Clara, CA, USA). The absorbance at O.D._660 nm_ was measured every 10 min.

### Motility assay

*C. crescentus* cultures grown in PYE were diluted to an O.D._660 nm_ of 0.4 and inoculated with sterile toothpicks on a PYE swarming plate (PYE + 0.25% agar). The plates were incubated at 30°C for 3 days, then imaged.

### Determination of cytoplasmic Cu content

*C. crescentus* cells were fixed for 20 min in 2% paraformaldehyde at 4°C and then washed three times with an ice-cold wash buffer (10 mM Tris-HCl pH 6.8, 100 μM EDTA). The cells were then incubated for 10 min with zwitterionic detergent (0.0125 % Zwittergent 3-14 detergent, 200 mM Tris-HCl pH 7.6), to disrupt the outer membrane but not the inner membrane. A final centrifugation at 14,500 x g for 15 min separated the spheroplasts (cytoplasm and inner membrane fraction) from the periplasm (and outer membrane fraction). The spheroplasts were lysed under 2.4 kbar by using a cell disrupter (Cell Disruption System, one shot model, Constant). Cells debris were removed by centrifugation at 10,000 x g for 10 min. and cell lysates were diluted in HNO_3_ 1 %. Samples were finally analyzed by inductively coupled plasma - optical emission spectrometry (ICP-OES) with an Optima 8000 ICP-OES from PerkinElmer. Cellular Cu concentration were calculated as described in (12). Briefly, 10 µl of the samples after fixation were diluted 1:100 in PBS and counted with a BD FACSVerseTM flow cytometer. Data were analyzed with the BD FACSuite V1.0.5 software. A ratio between the number of bacteria and the OD_660nm_ was determined. The combination of both flow cytometry and ICP-OES analyses allows to determine the number of Cu atoms in one spheroplast. Cu concentration was calculated using the following formula;

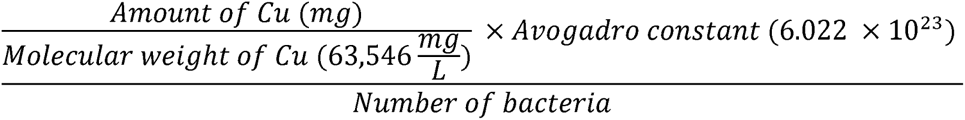

### Oxidative stress measurement with the CellROX Deep Red probe

An exponential phase culture of *C. crescentus* was synchronized to isolate the SW cells. The isolated SW cells were diluted to a final OD_660nm_ of 0.2 and incubated for 30 min at 30°C under moderate shaking with 5 μM of CellROX Deep Red probe (ThermoFisher) and appropriate concentrations of CuSO_4_ or H_2_O_2_. The bacteria were then washed three times with 20 mM Phosphate Buffer (12.5 mM Na_2_HPO_4_, 8 mM KH_2_PO_4_, pH 7) to remove the non-internalized probe. The bacteria were finally transferred into a black bottom 96-wells plate and the fluorescence was monitored at Em/Ex of 640/690 nm with a SpectraMax iD3 (Molecular Devices, San Jose, CA, USA). The relative fluorescence intensity was determined by defining 100% as the fluorescence from control condition.

### Oxidative stress measurement with the rxYFP biosensor

The SW cells were synchronized from an exponential phase culture overexpressing the rxYFP biosensor (47). The SW cells were diluted to a final OD_660nm_ of 0.4 in a black bottom 96-well plate and mixed with different concentrations of CuSO_4_ and H_2_O_2_. The rxYFP fluorescence was then monitored every minute for 20 min at Em/Ex of 510/556 nm with a SpectraMax iD3 (Molecular Devices, San Jose, CA, USA). The relative fluorescence intensity was determined by defining 100% as the initial measured fluorescence at time 0.

### Live Chemotaxis Imaging

The live chemotaxis imaging (LCI) assay was adapted from (12). Briefly, chemotaxis devices were made by casting solubilized 10:1 polydimethylsiloxane (PDMS Slygard 184, Dow Corning) in a small glass pot (d = 50 mm; h = 30 mm; glassware from Lenz laborglass instrument) in which coverslips were initially installed to mold the future bacterial chamber. The PDMS was left overnight, allowing degassing and curing. The different channels were drilled into the PDMS devices. A microscope slide and the PDMS cube were washed successively with acetone (only the slide), isopropanol, methanol and rinsed with milli-Q H_2_O. The materials were blown dry between each wash. The PDMS cube was sealed to the slide by pressure. Eight microliters of melted 1.5% agarose H_2_O with appropriate concentration of CuSO_4_, ZnSO_4_ or CdSO_4_ were loaded into the chamber through the external inlet channels to generate the plug. One hundred and fifty microliters of isolated SW cells were in turn injected into the bacterial chamber through the central inlet channel. Images were collected every minute for 25 min. in one focal plan (near the liquid surface) in the vicinity of the plug with a Nikon Eclipse Ti2-E inverted microscope equipped with a Nikon 20X/0.5 Ph2 objective and with a Hamamatsu C13440 digital camera. Quantitative analysis of the time-lapse images was performed with MicrobeJ (48). The number of bacteria in a same focal plane was determined at each time point; the number of bacteria at the beginning of the experiment is considered as 100%.

### Protein expression and purification

A BL21 pET28a-McpR_His_ exponential phase culture was incubated for 3 h under shacking at 37°C with 1 mM Isopropyl-β-D-thiogalactopyranoside (IPTG) and then harvested by centrifugation for 30 min at 4,000 x g at 4°C. McpR_His_ induction was verified by 12 % Tricine–sodium dodecyl sulfate–polyacrylamide gel electrophoresis (Tricine–SDS–PAGE), followed by Coomassie Blue staining and by Western-Blot using an anti-His antibody (MBL, PM032). To purify McpR_His_, the IPTG-induced culture was washed with binding buffer (20 mM Tris-HCl (pH 8.0), 500 mM NaCl, 10% glycerol, 10LJmM MgCl_2_) supplemented with 400 mg lysozyme (Sigma), complete EDTA-free protease cocktail inhibitor (Roche), 5 mg DNase I (Roche) and 0.4 g SDS and centrifuged for 15 min at 9,000 x g at 4°C. Bacterial cells were lysed at 2.4 kbar using a Cell disruptor (Cell Disruption System – one shot model, Constant). As McpR_His_ was mainly forming inclusion bodies, the pellet fraction was resuspended for 1 h under shaking at 4°C in the binding buffer containing 6 M guanidine. The solubilized inclusion bodies were centrifuged for 30 min at 9,000 x g at 4 °C. The collected supernatant was dialyzed overnight in a 0.1 M phosphate buffer (12.5 mM Na_2_HPO_4_, 8 mM KH_2_PO_4_, pH 8)) to remove guanidine, allowing the refolding of McpR. The same methodology was applied for the C75S and H99A mutants.

McpR protein concentration was determined (i) by measuring the absorbance of the sample at 280 nm with the SpectraMax iD3 (Molecular Devices, San Jose, CA, USA) or (ii) with a Bradford protein assay.

### Fluorescence quenching by Cu and other metal ions

Purified McpR at a concentration of 3.58 µM was incubated with different ratios of CuSO_4_, Tetrakis(acetonitrile)Cu(I) hexafluorophosphate, ZnSO_4_, CdSO_4_, MnSO_4_ and NiSO_4_ in black bottom 96-wells plates. The fluorescence emission spectra were recorded upon 280 nm excitation with the SpectraMax iD3.

### *K*d determination

The titration of McpR by Cu(I) or Cu(II) monitored by the protein intrinsic fluorescence intensity quenching was fitted to a single-site hyperbolic model by non-linear regression using the Scidavis 2.7 software to determine the affinity constants, according to the following equation:

> Fluorescence intensity = F0 + A*[Cu]/(Kd + [Cu]),

with F0 corresponding to the fluorescence intensity of the apo McpR, A to the total fluorescence amplitude change upon Cu binding and *K*d to the affinity dissociation constant. Dissociation constants are reported as the mean value ± standard deviation obtained from a minimum of three independent experiments performed on distinct protein productions.

### Identification of McpR oxidation by mass spectrometry

#### Protein digestion

McpR samples were treated using Filter-aided sample preparation (FASP) using the following protocol. To first wash the filter, 100 µl of formic acid 1 % were placed in each Millipore Microcon 30 MRCFOR030 Ultracel PL-30 before a 15 min centrifugation at 14,100 x g. For each sample, 40 µg of protein adjusted in 150 µl of 8 M urea buffer (8 M urea in 0.1 M Tris buffer at pH 8.5) were placed individually in a column and centrifuged for 15 min at 14,100 x g. The filtrate was discarded, and the columns were washed three times by adding 200 µl of urea buffer followed by a 15 min centrifugation at 14,100 x g. For the reduction step, 100 µl of dithiothreitol (DTT) were added and mixed for 1 min at 400 rpm with a thermomixer before a 15 min incubation at 24 °C. The samples were then centrifugated for 15 min at 14,100 x g, the filtrate was discarded, and the filter was washed by adding 100 µl of urea buffer before another 15 min centrifugation at 14,100 x g. An alkylation step was performed by adding 100 µl of iodoacetamide ((IAA), in urea buffer) in the column and mixing for 1 min at 400 rpm in the dark followed by a 20 min incubation in the dark and a 10 min centrifugation at 14,100 x g. To remove the excess of IAA, 100 µl of urea buffer were added and the samples were centrifuged for 15 min at 14,100 x g. To quench the rest of IAA, 100 µl of DTT were placed on the column, mixed for 1 min at 400 rpm and incubated for 15 min at 24 °C before a 10 min centrifugation at 14,100 x g. To remove the excess of DTT, 100 µl of urea buffer were placed on the column and centrifuged for 15 min at 14,100 x g. The filtrate was discarded, and the column was washed three times by adding 100 µl of 50 mM sodium bicarbonate buffer ((ABC), in ultrapure water) followed by a 10 min centrifugation at 14,100 x g. The remaining 100 µl were kept at the bottom of the column to avoid any evaporation in the column. The digestion process was performed by adding 80 µl of mass spectrometry grade trypsin (1/50 in ABC buffer) in the column and mixed for 1 min at 400 rpm followed by an overnight incubation at 24°C in a water saturated environment. The Microcon columns were placed in a 1.5 ml LoBind tube and centrifuged for 10 min at 14,100 x g. Fourty µl of ABC buffer were placed on the column before a 10 min centrifugation at 14,100 x g. Ten % Trifluoroacetic acid (TFA) in ultrapure water were added into the LoBind Tube to obtain 0.2 % TFA. The samples were dried in a SpeedVac up to 20 µl and transferred into an injection vial.

#### Mass spectrometry

The digest was analyzed using nano-LC-ESI-MS/MS tims TOF Pro (Bruker, Billerica, MA, USA) coupled with an UHPLC nanoElute (Bruker).

Peptides were separated by nanoUHPLC (nanoElute, Bruker) on a 75 μm ID, 25 cm C18 column with integrated CaptiveSpray insert (Aurora, ionopticks, Melbourne) at a flow rate of 400 nl/min, at 50°C. LC mobile phases A was water with 0.1% formic acid (v/v) and B ACN with 0.1% formic acid (v/v). Samples were loaded directly on the analytical column at a constant pressure of 800 bar. The digest (1 µl) was injected, and the organic content of the mobile phase was increased linearly from 2% B to 15 % in 36 min, from 15 % B to 25% in 19 min, from 25% B to 37 % in 5 min and from 37% B to 95% in 5 min. Data acquisition on the tims TOF Pro was performed using Hystar 5.1 and timsControl 2.0. tims TOF Pro data were acquired using 100 ms TIMS accumulation time, mobility (1/K0) range from 0.6 to 1.6 Vs/cm². Mass-spectrometric analysis were carried out using the parallel accumulation serial fragmentation (PASEF) acquisition method. One MS spectra followed by ten PASEF MSMS spectra per total cycle of 1.1 s.

Data analysis was performed using PEAKS Studio X Pro with ion mobility module (Bioinformatics Solutions Inc., Waterloo, ON). Protein identifications were conducted using PEAKS search engine with 15 ppm as parent mass error tolerance and 0.05 Da as fragment mass error tolerance. Carbamidomethylation was allowed as fixed modification, oxidation of methionine, histidine, tryptophan, and acetylation (N-term) as variable modification. Enzyme specificity was set to trypsin, and the maximum number of missed cleavages per peptide was set at three. The peak lists were searched against the *E. coli*-K12 (11002 sequences) and *Caulobacter* NA1000 (3863 sequences) from UNIREF 100. Peptide spectrum matches and protein identifications were normalized to less than 1.0% false discovery rate.

The identification results were processed with the PTM profile tool from PEAKS, which provides quantitative information of modified peptides compared with unmodified peptides for the modification sites of the protein across all MS samples (49,50).

### Circular dichroism

Purified McpR at a concentration of 8.95 µM was incubated with different CuSO_4_ and Tetrakis(acetonitrile)Cu(I) hexafluorophosphate concentrations. Measurements were performed with a MOS-500 spectropolarimeter in the far-UV region (190-260 nm), using a 1 mm pathlength cell. Four scans were averaged, and base lines were subtracted. Secondary structure analyses were performed on the circular dichroism data with the BeStSel server.

## Supporting information

This article contains supporting information.

## Acknowledgements

We would like to thank Jakob R. Winther for providing the rxYFP construct. We are also grateful to Valérie Charles and Carmela Aprile (CMI laboratory, NISM, UNamur) for the ICP measurements and to Virgile Neyman for technical support during the protein purification. Finally, we would like to acknowledge Jean-François Collet and Xavier De Bolle for critical reading of this manuscript and the URBM members and Johan Wouters (Chemistry Department, UNamur) for fruitful discussions.

## Funding and additional information

This work was supported by the University of Namur. G. L. was supported by the Belgian Fund for Industrial and Agricultural Research Associate (FRIA). Catherine Michaux and Eric Perpète thank the Belgian National Fund for Scientific Research for their respective associate and senior research associate positions.

## Data Availability

All data are contained within the manuscript and Supporting Information section. The raw data can be shared upon request to

---

**Figure S1:**
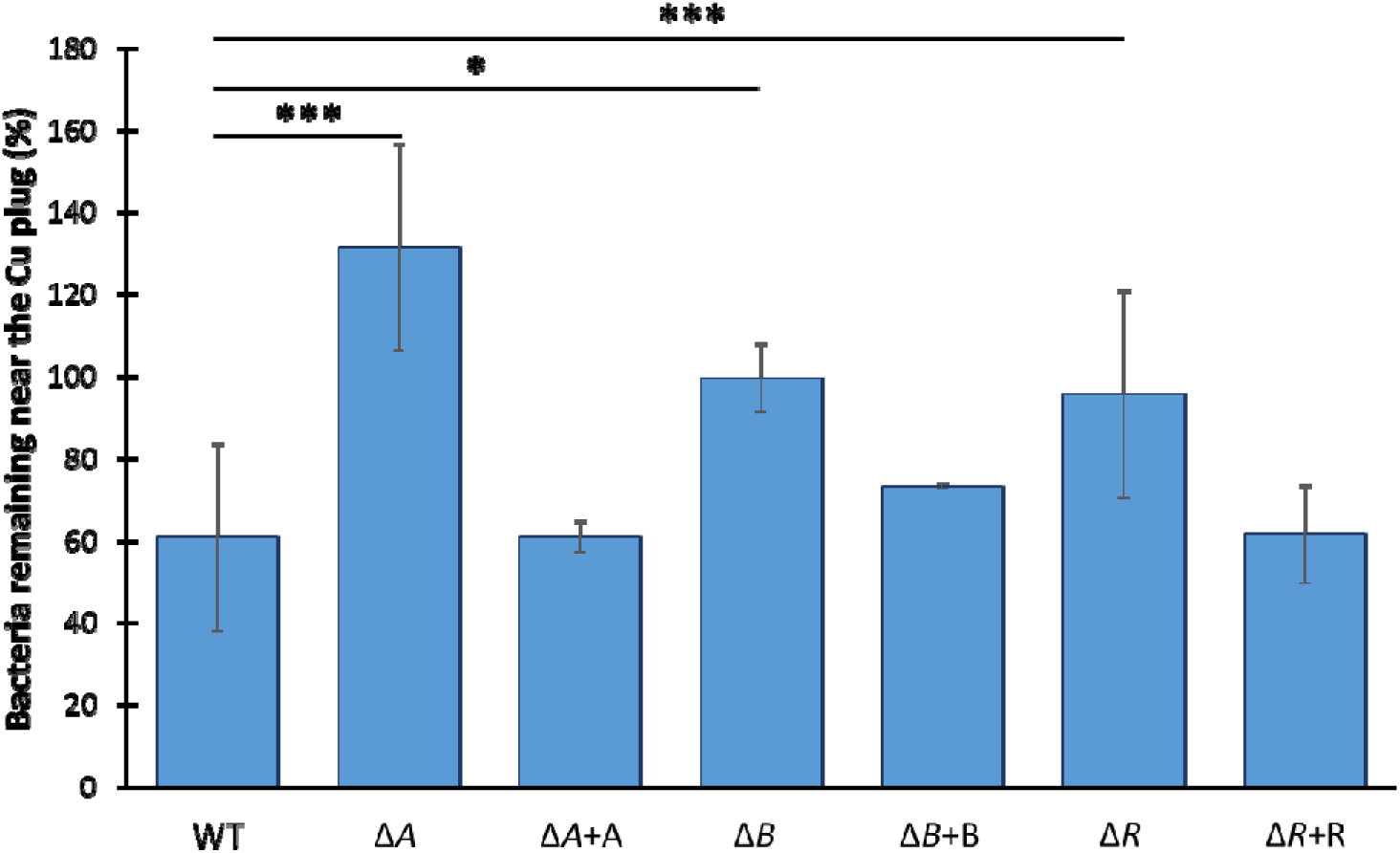
McpA, McpB and McpR are potentially involved in Cu chemotaxis. Percentage of WT, Δ*mcpA*, Δ*mcpA* overexpressing McpA, Δ*mcpB*, Δ*mcpB* overexpressing McpB, Δ*mcpR* and Δ*mcpR* overexpressing McpR SW cells remaining in the vicinity of the Cu plug after 25 min. Mean +/- s.d., at least 3 biological replicates. *p-values* were calculated using an ANOVA (* *p* < 0.05, ** *p* < 0.01, *** *p* < 0.001) (**Table S1**).

**Figure S2:**
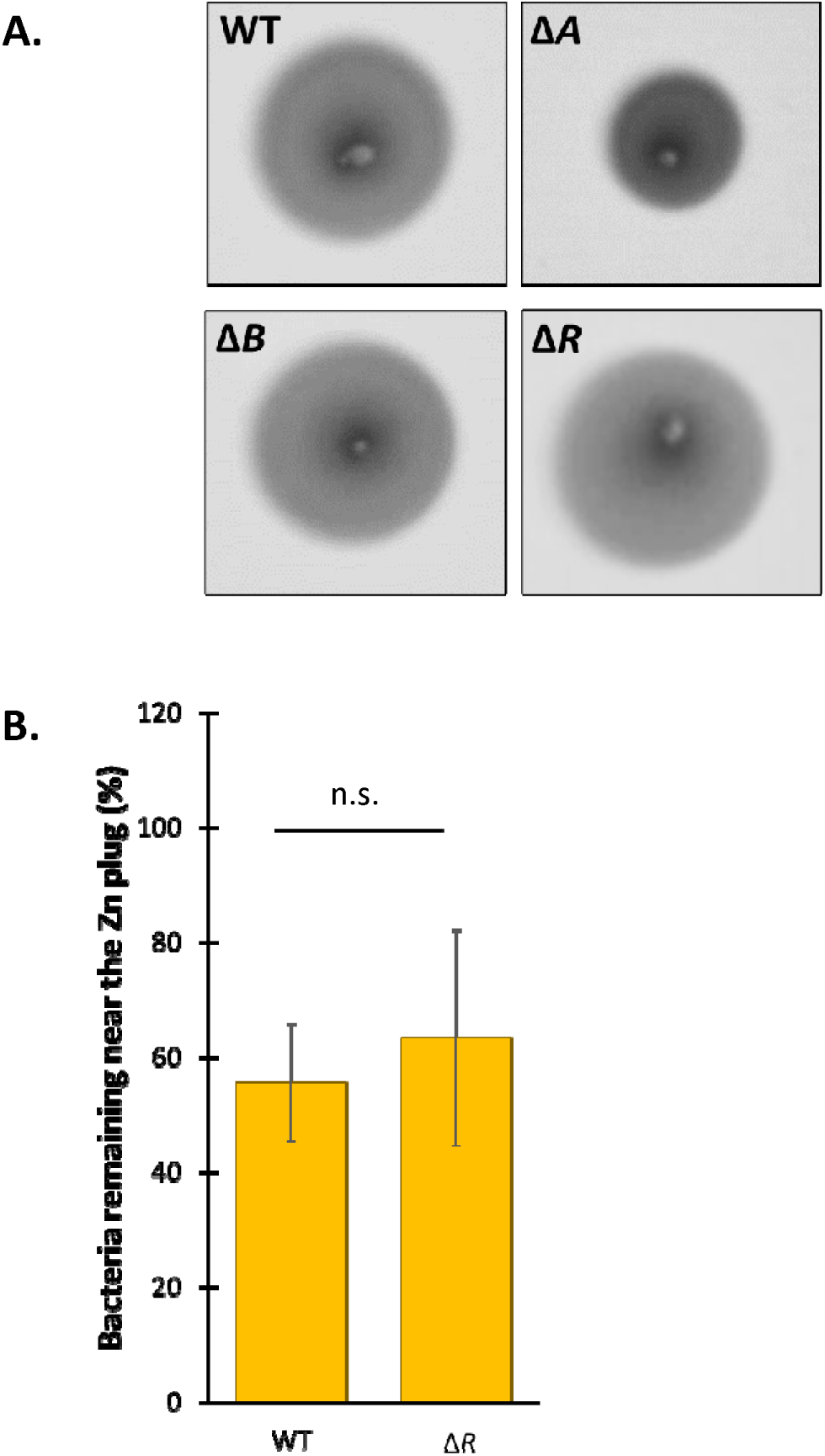
The chemotactic machinery is functional in the Δ*mcpR* mutant. **A.** Chemotaxis-driven swarming pattern of the WT, Δ*mcpA*, Δ*mcpB* and Δ*mcpR* strains in a Cu ions-free medium. **B.** Percentage of WT and Δ*mcpR* SW cells remaining in the vicinity of the Zn plug after 25 min. Mean +/- s.d., at least 3 biological replicates. *p-values* were calculated using a *t-test* (* *p* < 0.05, ** *p* < 0.01, *** *p* < 0.001) (**Table S1**).

**Figure S3:**
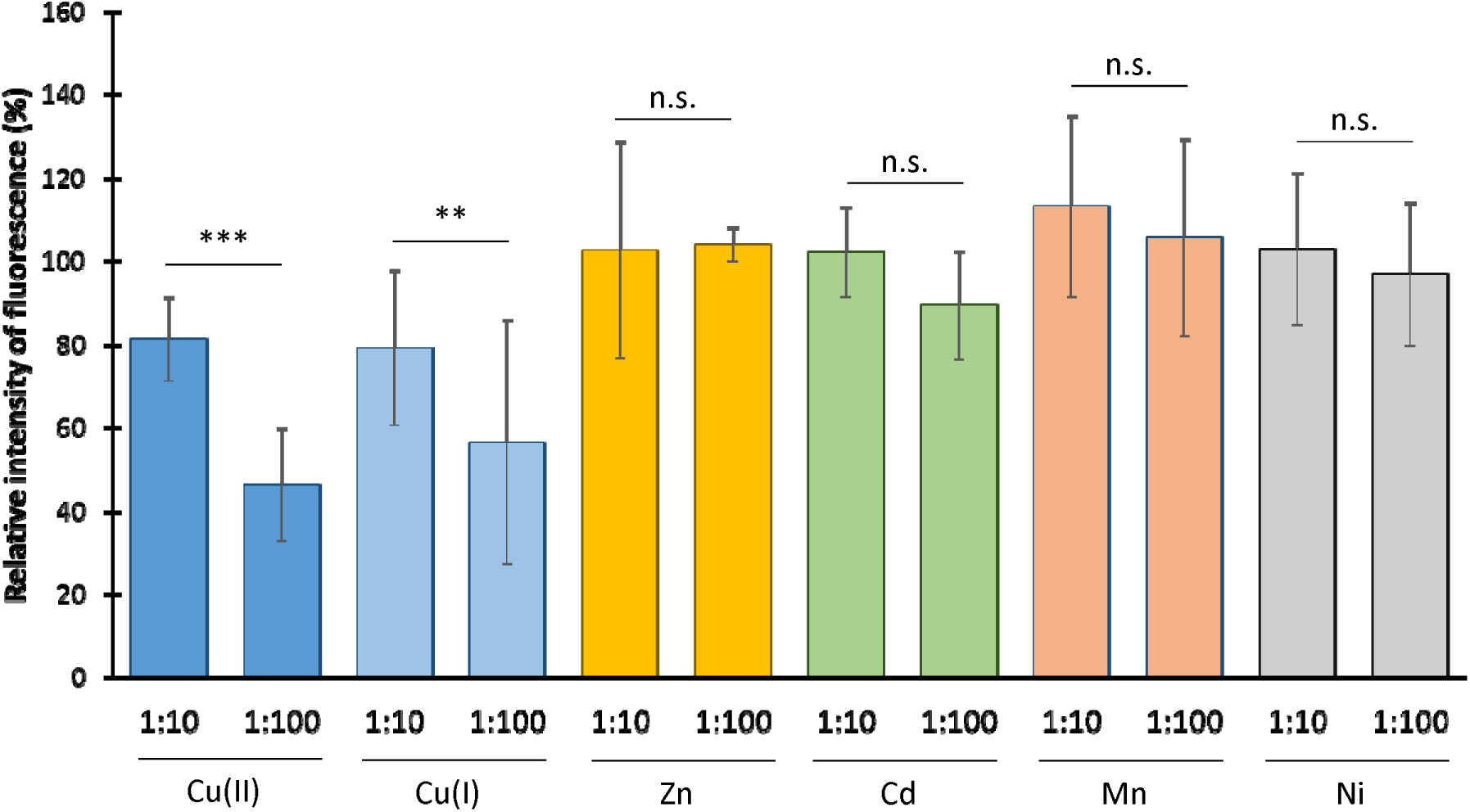
Zn, Cd, Mn and Ni do not directly bind to McpR. Relative intrinsic fluorescence of purified McpR (3.58 µM) incubated with ZnSO_4_, CdSO_4_, MnSO_4_ and NiSO_4_ at 1:10 and 1:100 ratios. Mean +/- s.d., at least 3 biological replicates. *p-values* were calculated using a t-test (* *p* < 0.05, ** *p* < 0.01, *** *p* < 0.001) (**Table S1**).

**Figure S4:**
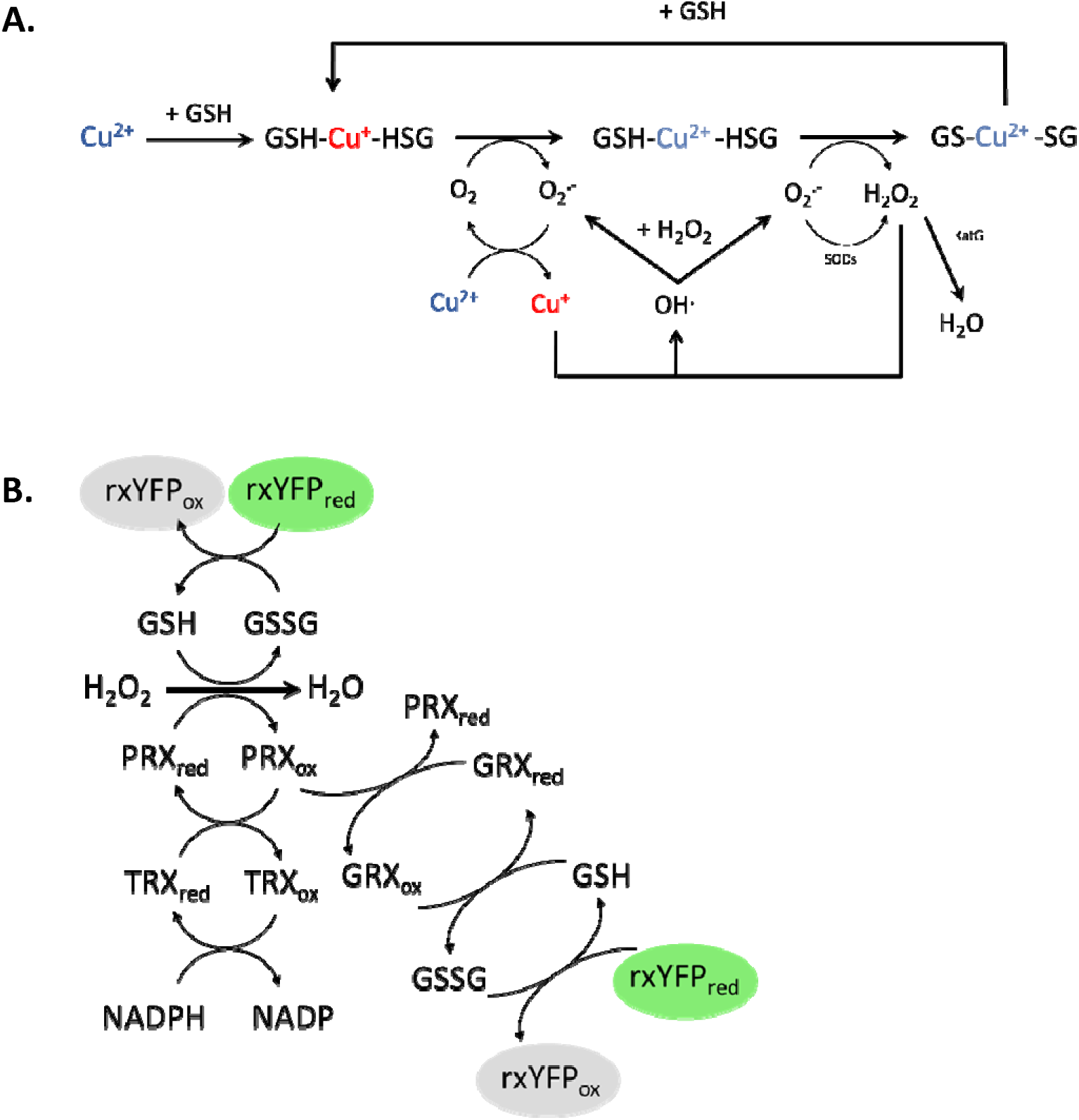
Potential pathways leading to ROS production by Cu. **A.** *In vitro* generation of Cu-induced ROS by Fenton-like reaction and Haber-Weiss cycles. The Cu-GSH complexes may also trigger ROS production (27, 28). The SODs and KatG are involved in O_2_•**-** and H_2_O_2_ detoxification, respectively **B.** The rxYFP biosensor is in equilibrium with the GSH pool. GSH is involved in different ways in ROS buffering. As a result, ROS production triggers an increase of the GS-SG pool, which will be partially reduced to GSH by using rxYFP as an electron donor. The resulting oxidized rxYFP will lose its fluorescence properties. GSH (glutathione), PRX (Peroxiredoxins), TRX (thioredoxin) and GRX (glutaredoxin).

**Figure S5:**
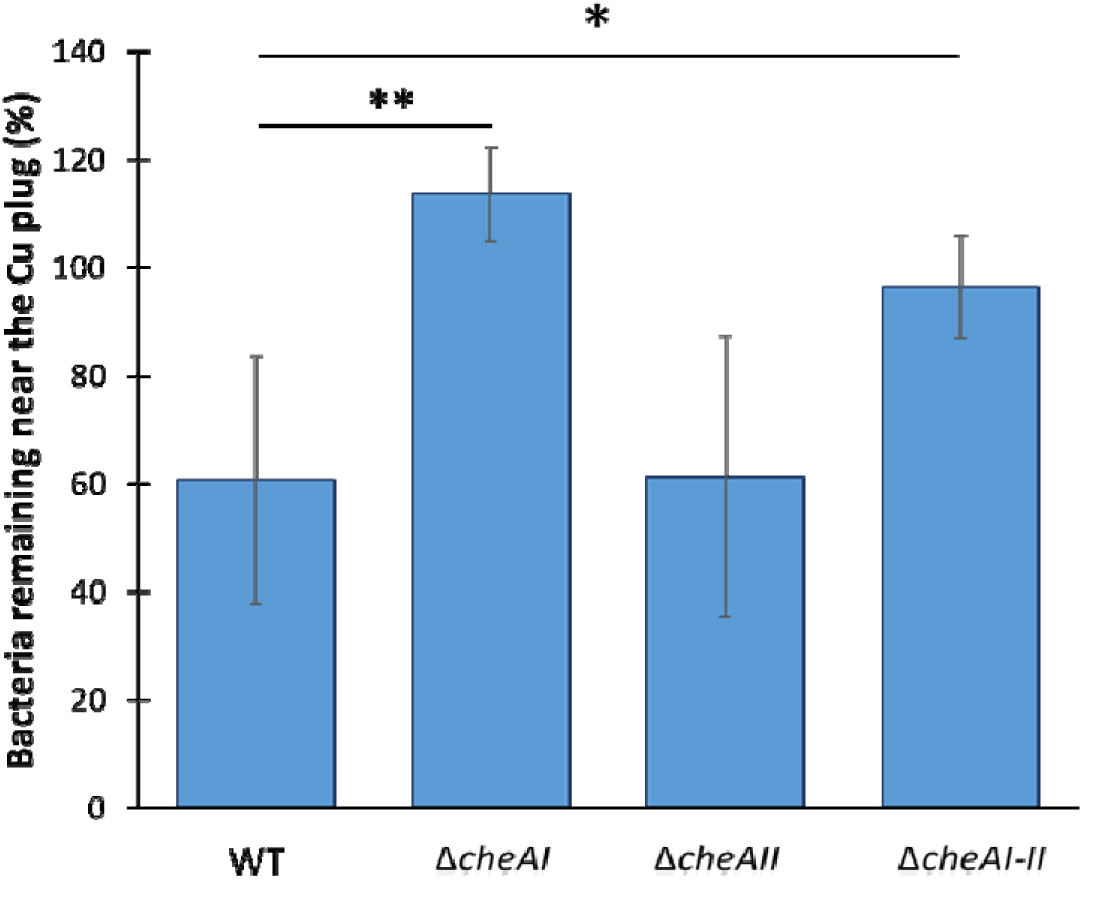
CheAI is the main histidine kinase involved in Cu chemotaxis. Percentage of WT, Δ*cheAI*, Δ*cheAII* and Δ*cheAI-II* SW cells remaining in the vicinity of the Cu plug after 25 min. Mean +/- s.d., at least 3 biological replicates. *p-values* were calculated using an ANOVA (* *p* < 0.05, ** *p* < 0.01, *** *p* < 0.001): WT vs. Δ*cheAI* = 0.0019; WT vs. Δ*cheAII* = >0.9999; WT vs. Δ*cheAI-II* = 0.0406.

**Figure S6:**
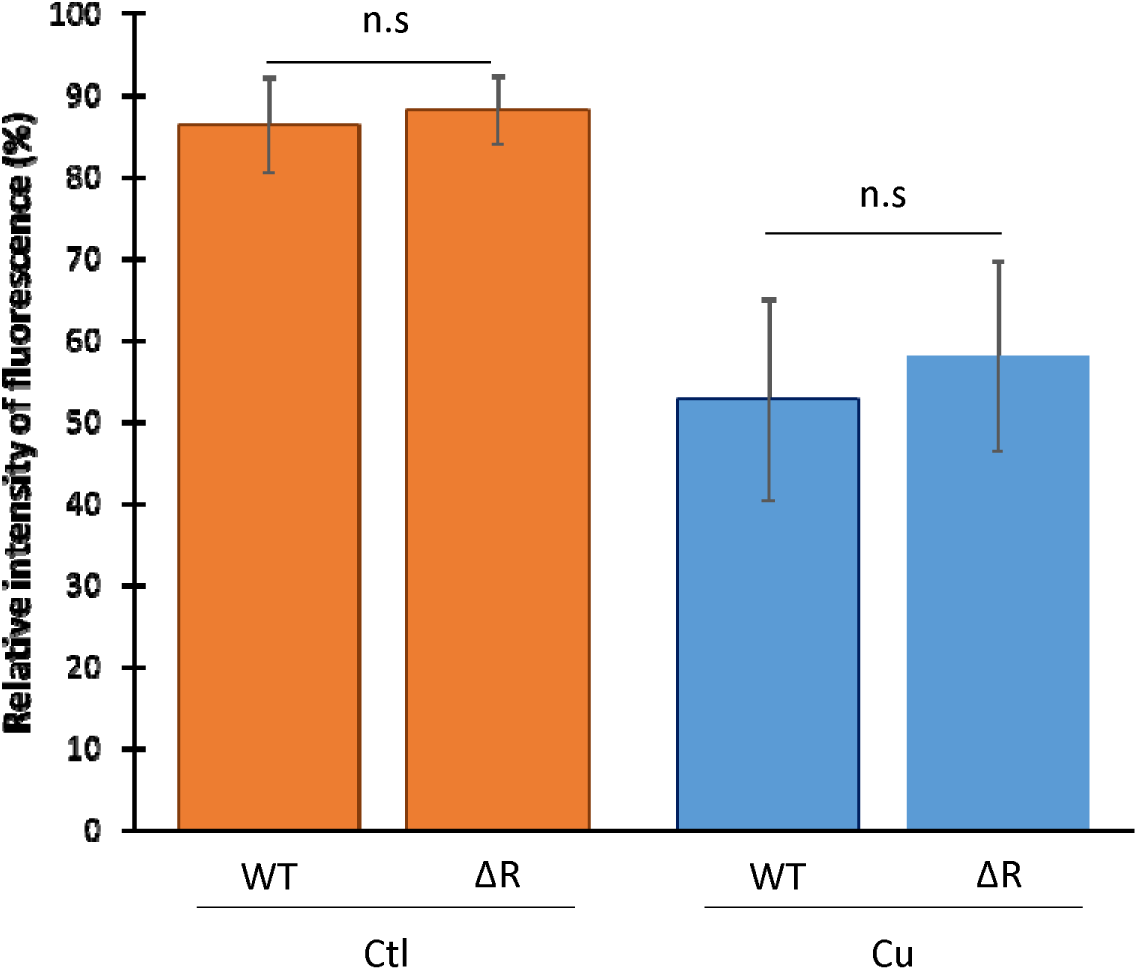
The Cu-induced oxidative stress is not affected in the Δ*mcpR* mutant. Relative intensity of fluorescence of the WT and Δ*mcpR* SW cells expressing the rxYFP biosensor and exposed to 175 µM Cu for 20 min. Mean +/- s.d., at least 3 biological replicates. *p-values* were calculated using a *t-test* (* *p* < 0.05, ** *p* < 0.01, *** *p* < 0.001) (**Table S1**).

**Figure S7:**
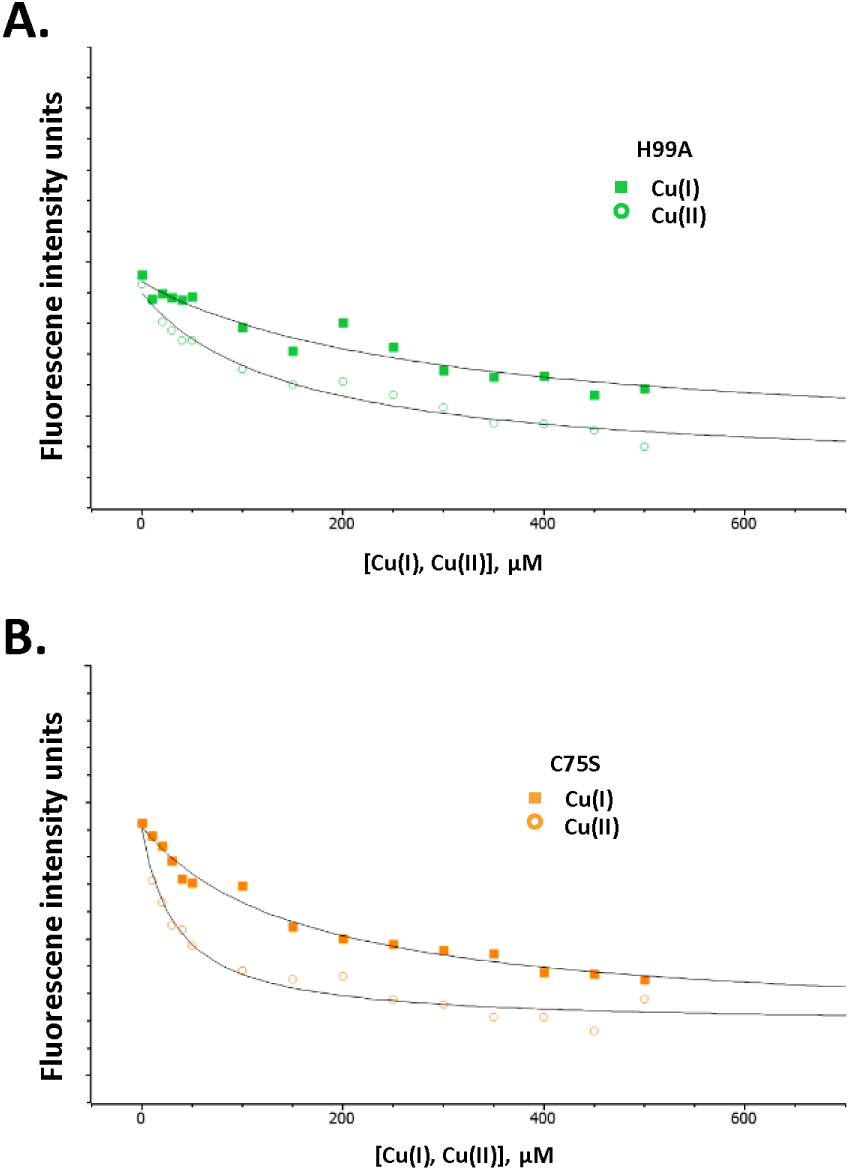
Fitting curves of the McpR^H99A^ and McpR_C75S_ variants. Representative H99A (**A.**) and C75S (**B.**) McpR titration by Cu(I) (squares) or Cu(II) (circles) monitored by protein intrinsic fluorescence intensity quenching, fitted to a single-site hyperbolic model by non-linear regression analysis (solid line). The fluorescence intensity axis scales correspond to the same amplitude for B and C.

**Figure S8:**
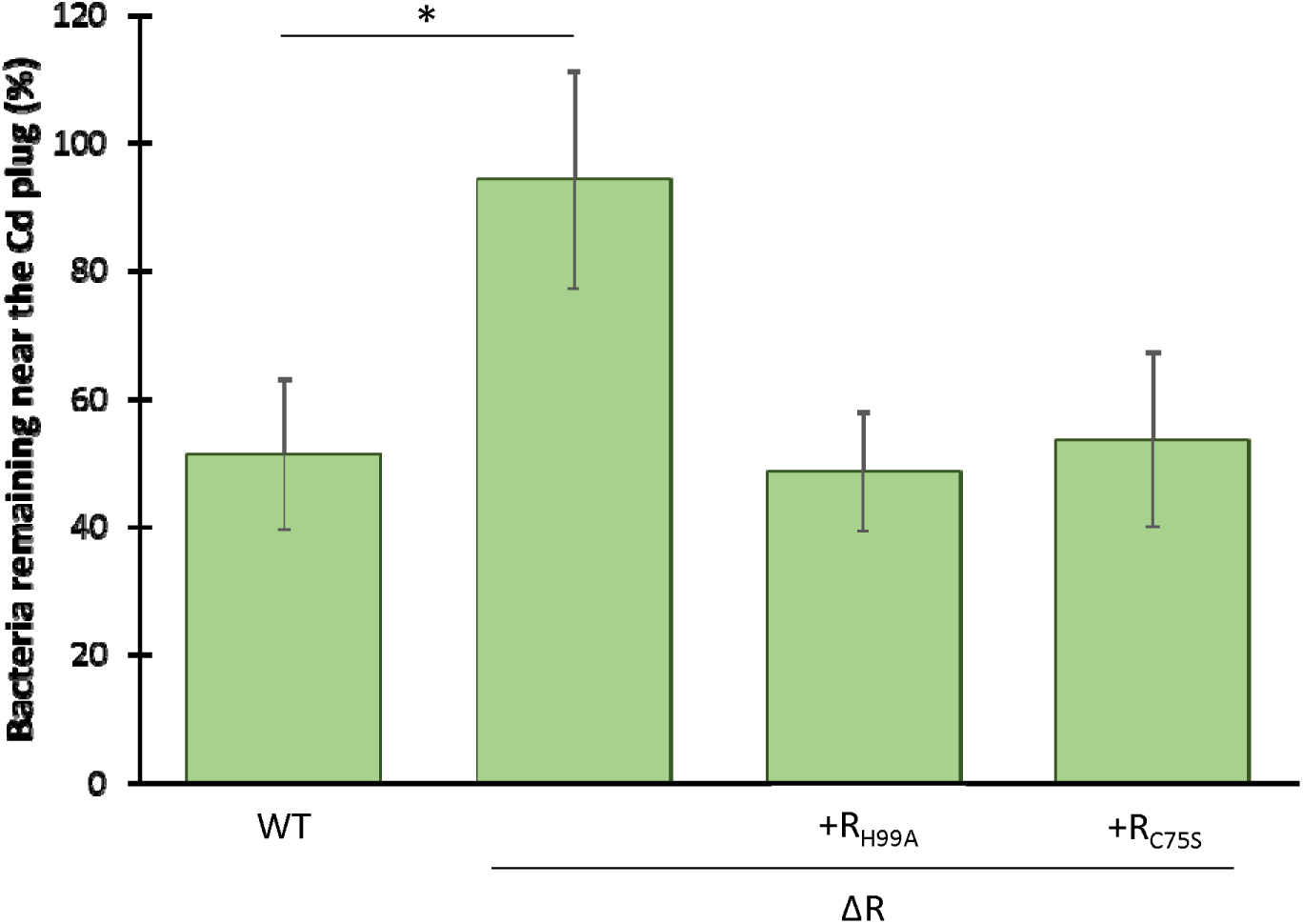
The McpR_C75S_ and McpR_H99A_ mutants are still functional. Percentage of WT, Δ*mcpR* and Δ*mcpR* overexpressing McpR, McpR_H99A_ or McpR_C75S_ SW cells remaining in the vicinity of the Cd plug after 25 min. Mean +/- s.d., at least 3 biological replicates. *p-values* were calculated using an ANOVA (* *p* < 0.05, ** *p* < 0.01, *** *p* < 0.001) (**Table S1**).

**Table S1:** *pvalues.* One-way ANOVA and t-test were performed using GraphPad Prism.

| **Figure 1A (ANOVA)** | **pvalue** | **summary** |
| --- | --- | --- |
| WT vs. ΔA | <0.0001 | **** |
| WT vs. ΔB | 0.0338 | * |
| WT vs. ΔC | 0.9993 | ns |
| WT vs. ΔD | 0.9992 | ns |
| WT vs. ΔE | >0.9999 | ns |
| WT vs. ΔF | 0.6094 | ns |
| WT vs. ΔG | 0.9991 | ns |
| WT vs. ΔH | 0.8952 | ns |
| WT vs. ΔI | 0.9689 | ns |
| WT vs. ΔJ | 0.999 | ns |
| WT vs. ΔK | 0.9991 | ns |
| WT vs. ΔL | >0.9999 | ns |
| WT vs. ΔM | 0.9796 | ns |
| WT vs. ΔN | 0.9993 | ns |
| WT vs. ΔO | 0.9996 | ns |
| WT vs. ΔP | 0.9996 | ns |
| WT vs. ΔQ | 0.9988 | ns |
| WT vs. ΔR | 0.0001 | *** |
| WT vs. ΔS | 0.9995 | ns |
| **Figure 1D (ANOVA)** | **pvalue** | **summary** |
| 50 μM Cu vs. 100 μM Cu | 0.0128 | * |
| 50 μM Cu vs. 175 μM Cu | 0.0011 | ** |
| 100 μM Cu vs. 175 μM Cu | 0.2469 | ns |
| **Figure 2A (ANOVA)** | **pvalue** | **summary** |
| Ctl vs. Cu | <0.0001 | **** |
| Ctl vs. H2O2 | 0.0007 | *** |
| **Figure 2B (ANOVA)** | **pvalue** | **summary** |
| Ctl vs. Cu | <0.0001 | **** |
| Ctl vs. H2O2 | <0.0001 | **** |
| **Figure 2C (ANOVA)** | **pvalue** | **summary** |
| Ctl vs. 20 μM | <0.0001 | **** |
| Ctl vs. 175 μM | <0.0001 | **** |
| Ctl vs. 300 μM | 0.0004 | *** |
| Ctl vs. 1.16 mM | <0.0001 | **** |
| **Figure 2D (ANOVA)** | **pvalue** | **summary** |
| WT vs. sodB+ | 0.0005 | *** |
| WT vs. katG+ | 0.001 | *** |
| **Figure 2F (ANOVA)** | **pvalue** | **summary** |
| WT vs. cheAI | 0.0007 | *** |
| WT vs. sodB+ | 0.0156 | * |
| WT vs. katG+ | 0.0045 | ** |
| **Figure 3C (ANOVA)** | **pvalue** | **summary** |
| WT vs. ΔR | 0.0001 | *** |
| WT vs. ΔR+R | >0.9999 | ns |
| WT vs. H99A | 0.2356 | ns |
| WT vs. C75S | 0.2503 | ns |
| ΔR+R vs. C75S | 0.0056 | ** |
| ΔR+R vs. H99A | 0.0053 | ** |

| **Figure S1 (ANOVA)** | **pvalue** | **summary** |
| --- | --- | --- |
| WT vs. ΔA | <0.0001 | **** |
| WT vs. ΔA+A | >0.9999 | ns |
| WT vs. ΔB | 0.0338 | * |
| WT vs. ΔB+B | >0.9999 | ns |
| WT vs. ΔR | 0.0001 | *** |
| WT vs. ΔR+R | 0.9954 | ns |
| **Figure S2 (t-test)** | **pvalue** | **summary** |
| WT vs. ΔR | 0.4878 | ns |
| **Figure S3 (t-test)** | **pvalue** | **summary** |
| Cu(II) 1:10 vs. Cu(II) 1:100 | 1.0372E-05 | *** |
| Cu(I) 1:10 vs. Cu(I) 1:100 | 0.00784501 | ** |
| Zn 1:10 vs. Zn 1:100 | 0.9479843 | ns |
| Cd 1:10 vs. Cd 1:100 | 0.59277248 | ns |
| Mn 1:10 vs. Mn 1:100 | 0.70207278 | ns |
| Ni 1:10 vs. Ni 1:100 | 0.58457574 | ns |
| **Figure S5 (t-test)** | **pvalue** | **summary** |
| WT Ctl vs. R Ctl | 0.9651 | ns |
| WT Cu vs. R Cu | 0.6783 | ns |
| **Figure S7 (ANOVA)** | **pvalue** | **summary** |
| WT vs. ΔR | 0.0171 | * |
| WT vs. H99A | 0.8499 | ns |
| WT vs. C75S | 0.9949 | ns |

**Table S2:**
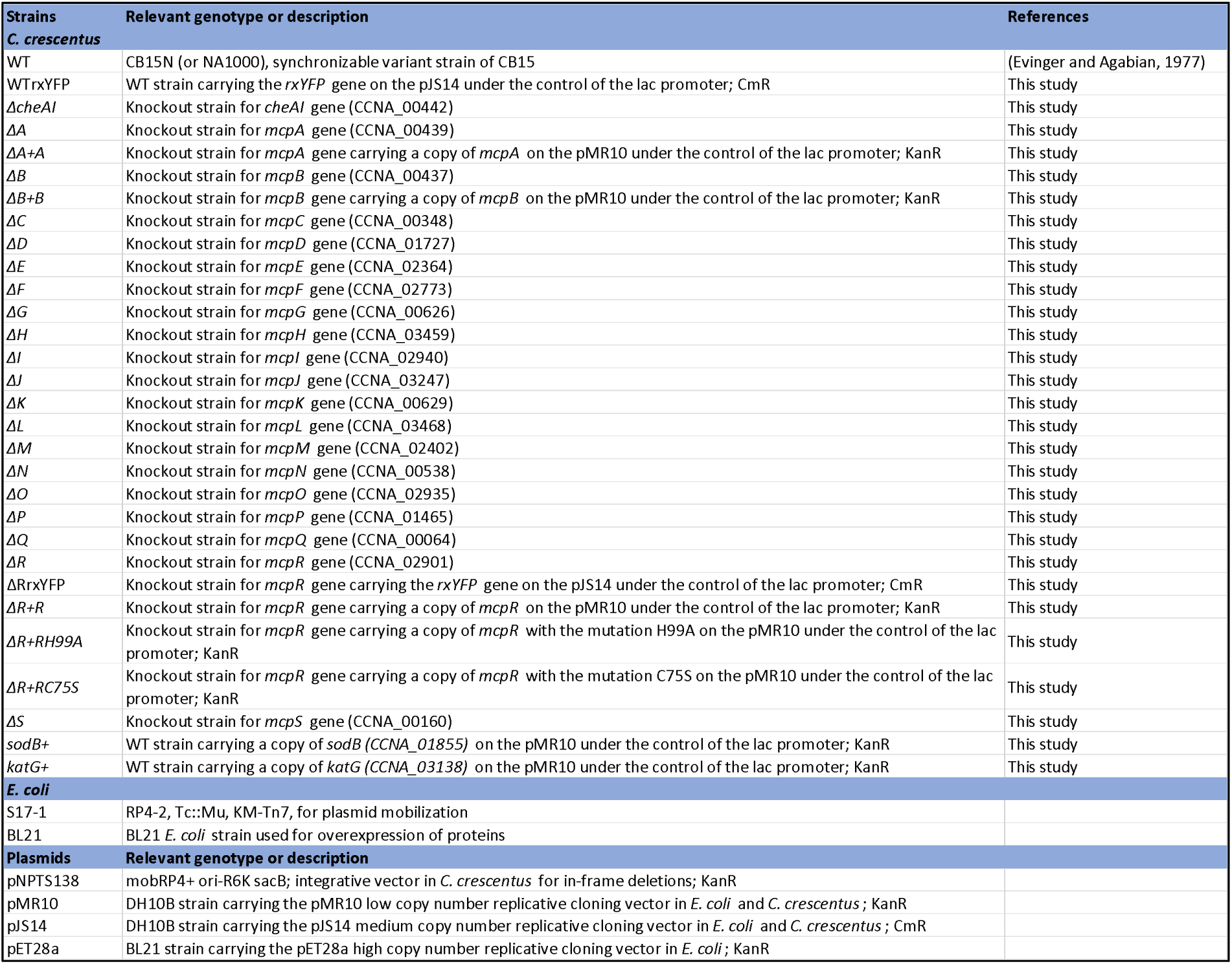
Strains, primers and plasmids (1/4)

**Table S2:**
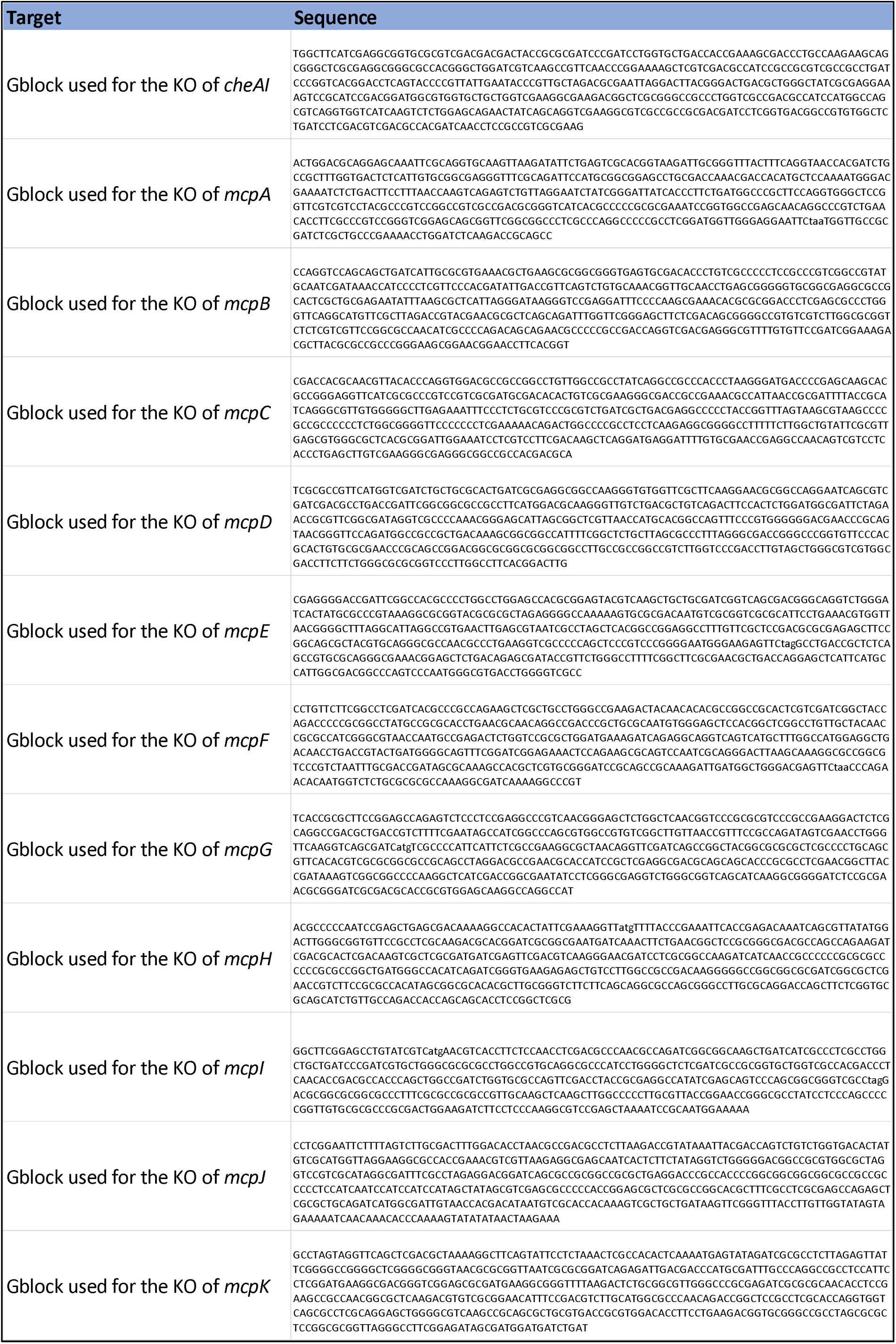
Strains, primers and plasmids (2/4)

**Table S2:**
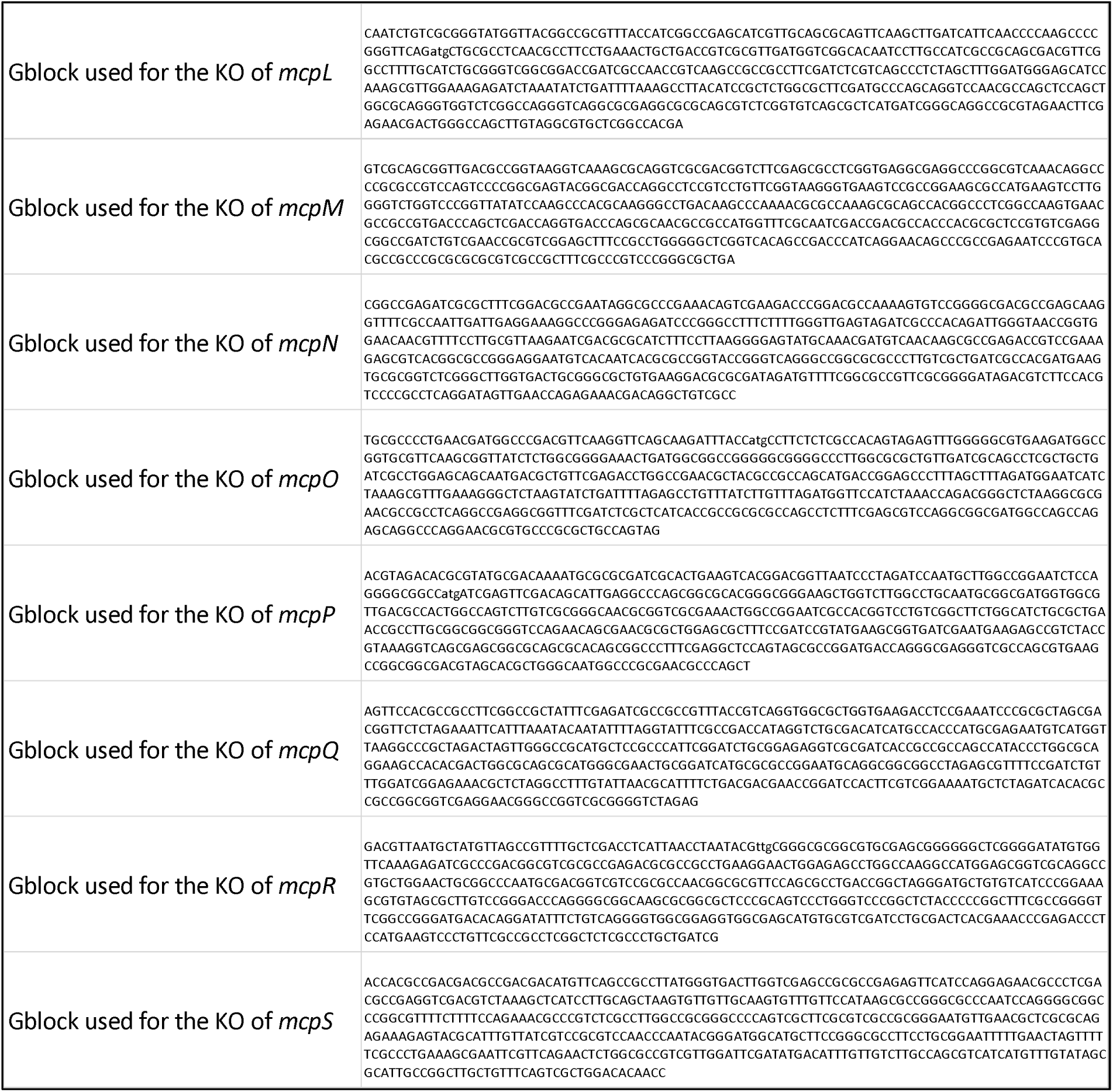
Strains, primers and plasmids (3/4)

**Table S2:**
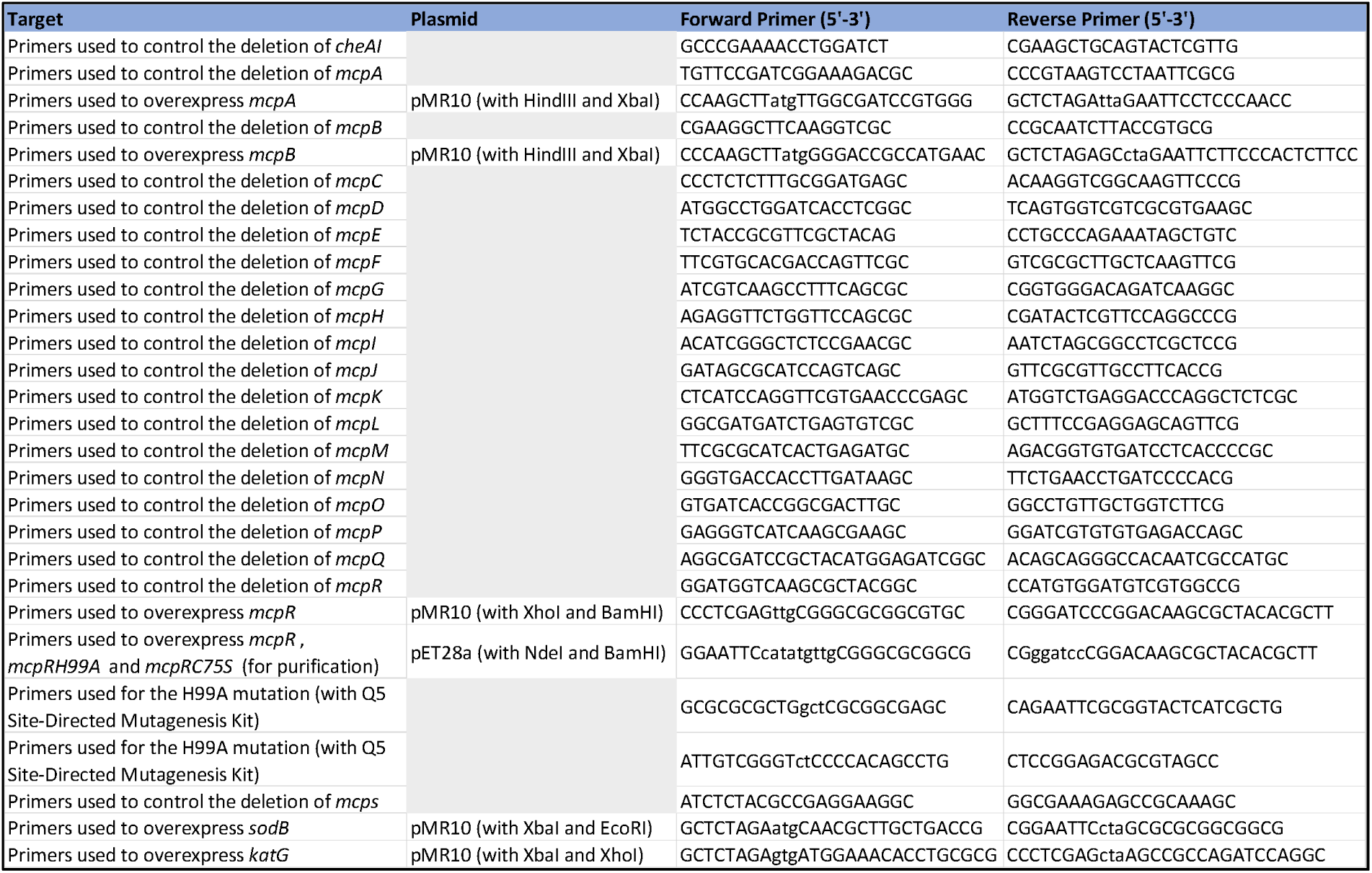
Strains, primers and plasmids (4/4)

