## Supporting Information for "A cytoplasmic chemoreceptor and Reactive Oxygen Species mediate bacterial chemotaxis to copper"

*Caulobacter crescentus* genome is predicted to encode two chemotaxis clusters including genes coding for CheA histidines kinases, CheY response regulators, CheW adaptors, CheB methylesterases, CheR methytransferases, McpA and McpK and other uncharacterized Che proteins. Most of the studies conducted in *C. crescentus* so far focused on the major chemotaxis cluster (cluster 1) (1-4) while the second alternate chemotaxis cluster (cluster 2) was only recently investigated (1). These studies concluded that the cluster 1 is essential for chemotaxis, while cluster 1 and cluster 2 are involved in biofilm formation and holdfast production (1, 2). Consistent with these observations, CheAI (cluster 1 CheA) turns out to be the main histidine kinase involved in chemotaxis while the exact function of CheAII (cluster 2 CheA) remains unknown^1^. In order to determine which CheA is involved in Cu-chemotaxis, we constructed the Δ*cheAI*, Δ*cheAII* and Δ*cheAI-II* mutants and assessed their chemotactic behavior to Cu by LCI (**Fig. S5**). The Δ*cheAI* mutant completely lost its chemotactic response to Cu, while the Δ*cheAII* mutant displayed a WT phenotype. Accordingly, the Δ*cheAI-II* double mutant showed a similar behavior to the Δ*cheAI* mutant. These data confirm that the CheAI histidine kinase is the main histidine kinase involved in Cu chemotaxis.
